# CoMM-S^2^: a collaborative mixed model using summary statistics in transcriptome-wide association studies

**DOI:** 10.1101/652263

**Authors:** Yi Yang, Xingjie Shi, Yuling Jiao, Jian Huang, Min Chen, Xiang Zhou, Lei Sun, Xinyi Lin, Can Yang, Jin Liu

**Affiliations:** School of Statistics and Management, Shanghai University of Finance and Economics, Shanghai, China; Centre for Quantitative Medicine, Program in Health Services and Systems Research, Duke-NUS Medical School, Singapore 169857; Department of Statistics, Nanjing University of Finance and Economics, Nanjing, China; School of Statistics and Mathematics, Zhongnan University of Economics and Law, Wuhan, China; Department of Applied Mathematics, Hong Kong Polytechnics University, Hong Kong, China; Academy of Mathematics and Systems Science, The Chinese Academy of Sciences, Beijing, China; Department of Biostatistics, University of Michigan, Ann Arbor, MI 48109, USA; Cardiovascular and Metabolic Disorders Program, Duke-NUS Medical School, Singapore; Singapore Clinical Research Institute, Singapore; Singapore Institute for Clinical Sciences, A*STAR, Singapore; Department of Mathematics, Hong Kong University of Science and Technology, Hong Kong, China

## Abstract

**Motivation:** Although genome-wide association studies (GWAS) have deepened our understanding of the genetic architecture of complex traits, the mechanistic links that underlie how genetic variants cause complex traits remains elusive. To advance our understanding of the underlying mechanistic links, various consortia have collected a vast volume of genomic data that enable us to investigate the role that genetic variants play in gene expression regulation. Recently, a collaborative mixed model (CoMM) [42] was proposed to jointly interrogate genome on complex traits by integrating both the GWAS dataset and the expression quantitative trait loci (eQTL) dataset. Although CoMM is a powerful approach that leverages regulatory information while accounting for the uncertainty in using an eQTL dataset, it requires individual-level GWAS data and cannot fully make use of widely available GWAS summary statistics. Therefore, statistically efficient methods that leverages transcriptome information using only summary statistics information from GWAS data are required.

**Results:** In this study, we propose a novel probabilistic model, CoMM-S^2^, to examine the mechanistic role that genetic variants play, by using only GWAS summary statistics instead of individual-level GWAS data. Similar to CoMM which uses individual-level GWAS data, CoMM-S^2^ combines two models: the first model examines the relationship between gene expression and genotype, while the second model examines the relationship between the phenotype and the predicted gene expression from the first model. Distinct from CoMM, CoMM-S^2^ requires only GWAS summary statistics. Using both simulation studies and real data analysis, we demonstrate that even though CoMM-S^2^ utilizes GWAS summary statistics, it has comparable performance as CoMM, which uses individual-level GWAS data.

**Availability and implementation:** The implement of CoMM-S^2^ is included in the *CoMM* package that can be downloaded from https://github.com/gordonliu810822/CoMM.

**Supplementary information:** Supplementary data are available at *Bioinformatics* online.

## 1 Introduction

Over the last decade, genome-wide association studies (GWAS) have achieved remarkable success in identifying genetic susceptibility variants for a variety of complex traits/diseases [39]. However, the biology of how genetic variants affect complex traits remains unclear. Recent expression quantitative trait loci (eQTL) studies indicate that regulatory information play an important role in mediating the complex traits/diseases [26]. Measured comprehensive cellular traits can serve as reference data and provide investigators with an avenue to examine the role that genetic variants play in gene expression regulation. For example, the Genotype-Tissue Expression (GTEx) Project [23] has provided DNA sequencing data (about 12.5 million variants) from 449 individuals and collected gene-expression measurements of 44 tissues from these individuals; the number of subjects increases to 620 in over 48 tissues in the recent V7 release. Although the sample sizes of these reference datasets are limited, they provide an important avenue for one to study how genetic variants regulate human gene expression in different tissues.

In the absence of identical cohorts in eQTL and GWAS datasets, various authors have proposed statistical methods that allow one to leverage regulatory information on the cellular mechanisms in a GWAS analysis. These methods can be broadly grouped into two categories. The first group consists of methods that require the use of individual-level GWAS data, and include methods such as PrediXcan [10] and CoMM [42]. Because methods in this category require the availability of all individual-level genotype and phenotype data, their application can be complicated by restrictions on data sharing and storage. In contrast to the first group of methods that utilize individual-level data, the second group of methods uses GWAS summary statistics; for example one could apply the second group of methods to GWAS results that are publicly available from GWAS repositories, such as the NHGRI-EBI GWAS Catalog [4]. Examples of these methods include TWAS [13], S-PrediXcan [1], and UTMOST [17]. Among these methods, TWAS and S-PrediXcan use transcriptome data from a single tissue while UTMOST can be applied to cross-tissue analysis. Generally, TWAS-type methods (PrediXcan, S-PrediXcan, TWAS and UTMOST) proceed with three steps. First, the expression reference panel is used to fit predictive models for each gene using genetic variants in the vicinity of a gene. Next, levels of gene expression for the individuals in the GWAS data are predicted using these models. Finally, associations between the predicted expression levels and the complex trait are examined by simple linear regressions. Consequently, TWAS-type methods do no account for the uncertainty associated with the first step. In contrast, CoMM accounts for the uncertainty by combining the three steps in a unified probabilistic framework.

Compared with methods using individual-level GWAS data, methods using GWAS summary statistics face an additional difficulty: the summary statistics do not contain any information of linkage disequilibrium (LD), which plays an important role in prioritizing variants in GWAS. TWAS used an imputation method to impute the expression-trait association statistics directly from GWAS summary statistics [13] while S-PrediXcan derived a test statistic using pre-calculated weights to expression and a reference panel to estimate correlation (LD) among cis-variants. To make use of summary statistics rigorously, it is important to develop a probabilistic model. [16] first proposed an approximated distribution for z-scores in CAVIAR. Later, [45] formalized this distribution by introducing a regression with summary statistics (RSS) likelihood in a Bayesian framework and they further showed that the difference between the RSS log-likelihood and the one from individual-level data was constant. Although the approximated RSS-type distribution has been extended in several works including RSS-E [46] and REMI [18], both of these works including RSS are designed for one-sample studies. In the analysis using two different samples, such as PrediXcan, TWAS and CoMM, the questions become how to combine a RSS distribution for GWAS summary statistics with that for eQTL data.

To overcome the limitation of not accounting for uncertainty and further extend CoMM using GWAS summary statistics, we propose a probabilistic model, a <u>Co</u>llaborative <u>M</u>ixed <u>M</u>odels for GWAS <u>s</u>ummary <u>s</u>tatistics – CoMM-S^2^. Unlike TWAS and S-PrediXcan, our method accounts for the uncertainty in the ‘imputed’ gene expression. The key idea is to build a joint probabilistic model for GWAS summary statistics and individual-level eQTL data, and use the 1KG data as a reference panel to estimate LD. We has also developed an efficient variational Bayesian expectation-maximization accelerated using parameter expansion (PX-VBEM), where the calibrated evidence lower bound is used to conduct likelihood ratio tests for genome-wide gene associations with complex traits/diseases. We illustrate the performance of CoMM-S^2^ with extensive simulation studies and real data applications of 10 traits in NFBC1966 dataset and summary statistics from 14 traits/diseases. The results demonstrate that CoMM-S^2^ performs better than competing methods.

## 2 Methods

### 2.1 Notation

Suppose that we have an individual-level eQTL dataset 𝒟_1_ = {**Y, W**_1_} that consists of *n*_1_ samples, *g* genes and *m* genetic variants, and where 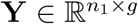 is a matrix of gene expression and 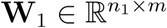 is a genotype matrix. In addition, we have the GWAS summary statistics 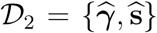, where 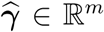 and 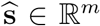 are the effect sizes and standard errors from the single-variant analysis for all genetic variants. We further assume that the individual-level GWAS data corresponding to 𝒟_2_ has the phenotype vector 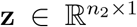 and the centered genotype matrix 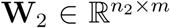. The sample sizes of 𝒟_1_ and 𝒟_2_ are generally distinct, with 𝒟_1_ having a smaller sample size (≈ 10^2^) compared to the sample size of 𝒟_2_ (≈ 10^4^ ∼ 5×10^5^). We will examine gene expression levels for each gene individually. Let **y**_*j*_, the *j*-th column of **Y**, be the gene expression level of the *j*-th gene and let **W**_1*j*_ be a genotype matrix containing its nearby genetic variants (within either 50 kb upstream of the transcription start site or 50 kb downstream of the transcription end site, in this study), respectively. We standardize the genotype data 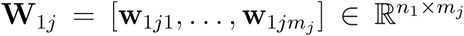 to mean zero and unit variance. Correspondingly, summary statistics for genetic variants using the centered genotype (**W**_2*j*_) within the *j*-th gene is 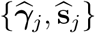, where 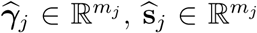, and *m*_*j*_ is the number of variants corresponding to the *j*-th gene. Denote 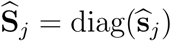 a diagonal matrix for the *j*-th gene and 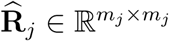 the estimated correlation among genetic variants within the *j*-th gene.

### 2.2 Model

We first model the relationship in the eQTL data using linear regression

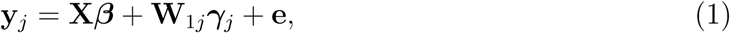

where 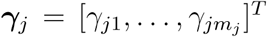 is an *m*_*j*_ × 1 vector of genetic effects on the gene expression, 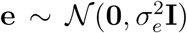 is an *n*_1_ × 1 vector of independent random noises for the gene expression levels, 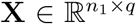 are the design matrix for covariates including intercept, and ***β*** is a *q* × 1 vector of the corresponding effect sizes for covariates. Similar to CoMM [42], the relationship between the phenotype **z** and genotype **W**_2*j*_ nearby the *j*-th gene can be modeled as

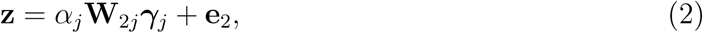

where 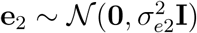 is an *n*_2_ ×1 vector of independent error associated with the phenotype. Here, *α*_*j*_ represents the effects of gene expression of gene *j* on the phenotype, due to genotype. Assume that individual-level data {**z, W**_2_} is inaccessible, but the summary statistics 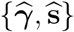 from the univariate linear regression are available. It can be shown that the distribution of 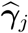 can be approximated by [45]

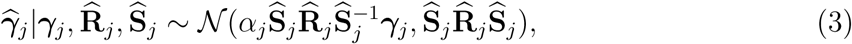

provided that sample size *n*_2_ to generate these summary statistics is large and the trait is highly polygenic (*i.e.*, the squared correlation coefficient between the trait and each genetic variant is close to zero). We further assume the prior distribution for ***γ***_*j*_ is a Gaussian,

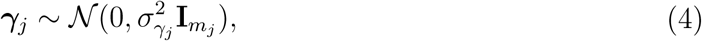

which is widely used in genetics [44]. Taking ***γ***_*j*_ as the latent variable, the complete data likelihood can be written as

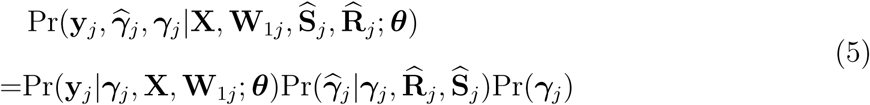

where 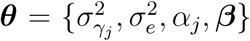 is the collection of model parameters. By integrating out latent variable ***γ***_*j*_, the marginal likelihood is

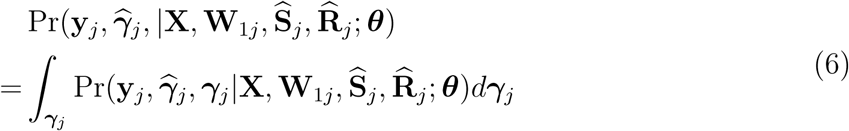

### 2.3 Algorithm

We require a computationally efficient algorithm that is capable of fitting model (5) when the signal-noise-ratio is low. A standard expectation-maximization (EM) algorithm in not ideal for this purpose due to the slow convergence; a Newton-Raphson algorithm is also not ideal because it can be unstable because of the non-negative constraint on variance components. Additionally, a standard EM algorithm involves the inversion of 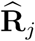, which may cause numerical failure as 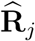 is estimated from a small reference panel. Therefore, we develop a variational Bayesian (VB) EM algorithm [3] accelerated by parameter expansion [22], namely, PX-VBEM. First, the original model (1) can be expanded as follows

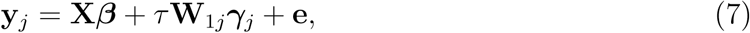

where *τ* ∈ℝ is the expanded parameter, the likelihood for summary statistics (3) and the prior remain the same, and model parameters become 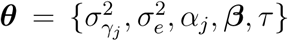. Next, we sketch the variational Bayesian EM algorithm for the expanded model (7). Given a variational posterior distribution *q*(***γ***_*j*_), it is easy to verify that the marginal likelihood can be decomposed into two components, the evidence lower bound (ELBO) and the KL divergence,

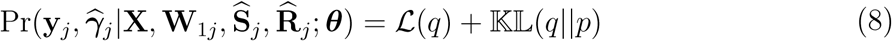

where

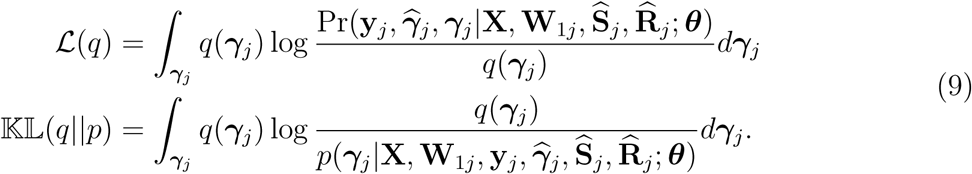

Note that ℒ(*q*) is the ELBO of the marginal likelihood, and 𝕂𝕃(*q‖p*) is Kullback-Leibler (KL) divergence between two distributions and satisfies 𝕂𝕃(*q‖p*) *≥* 0, with the equality holding if, and only if, the variational posterior probability (*q*) and the true posterior probability (*p*) are equal. Similar to the EM algorithm, we can maximize the ELBO ℒ(*q*) by optimizing with respect to *q* that is equivalent to minimizing the KL divergence [2]. To make the evaluation of the lower bound computationally efficient, we use the mean-field theory [27] and assume that *q*(***γ***_*j*_) can be factorized as

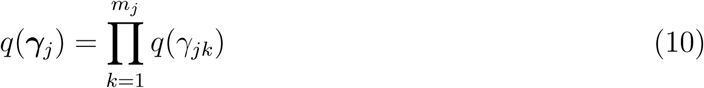

This is the only assumption that we make using variational inference. This factorization (10) is used as an approximation for the posterior distribution 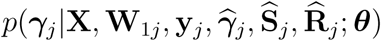. In the VB E-step, given the hidden variables *γ*_*ji*_, *i ≠ k*, the terms with *γ*_*jk*_ have a quadratic form, where *i* and *k* are indices for the *i*-th and the *k*-th genetic variants, respectively. Thus, the variational posterior distribution of *γ*_*jk*_ is a Gaussian distribution 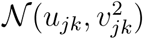. The details of derivation for the updating formula of mean 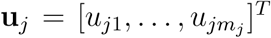 and standard deviation 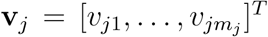, and the ELBO ℒ(*q*) of the marginal likelihood (9) at the old parameters ***θ***^*old*^ can be found in the supplementary document. In the VB M-step, we obtain the new updates for all parameters ***θ*** by setting the derivative of ELBO to zero. The resulting updating equations for parameters are

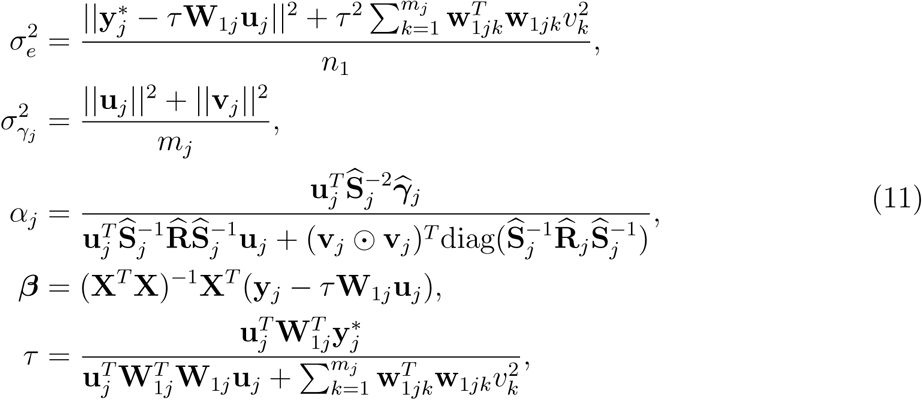

where 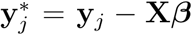 is the vector of residuals after removing the fixed effects, *0* denotes the element-wise multiplication of two vectors, and diag(**A**) denotes a vector whose elements are the diagonal entries of the square matrix **A**. The corresponding PX-VBEM algorithm is summarized as Algorithm 1 in the supplementary document.

### 2.4 Reference panel

The CoMM-S^2^ uses marginal effect sizes and their standard errors to construct probabilistic modeling for summary statistics from GWAS. Using summary-level data, we do not have any information for correlations among SNPs (*i.e.*, LD, denoted as **R**_*j*_). Here, we choose to use 1KG samples as a reference panel. We first calculate the empirical correlation matrix 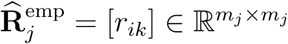 with 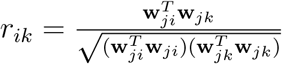, where **w**_*jk*_ is the genotype vector for the *k*-th genetic variant within the *j*-th gene. To make the estimated correlation matrix positive definite, we applied a simple shrinkage estimator [32] to obtain 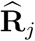 as 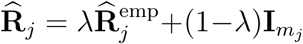, where *Λ ∈* [0, 1] is the shrinkage intensity. Note that the shrinkage correlation matrix is the combination of the two extremes, the empirical correlation matrix 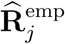 and the identity matrix **I**. It is easy to recognize that the shrinkage correlation matrix can recover the original empirical correlation matrix 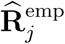 when *Λ* = 1 or identity matrix **I** when *Λ* = 0. In addition, we have tested with different *Λ ∈* [0.8, 0.95] for CoMM-S^2^ and its results are quite robust.

## 3. Statistical Inference

### 3.1 Evaluate association between a a complex trait/disease and a gene

We propose the following statistical test to formally examine the association between a a complex trait/disease and a gene:

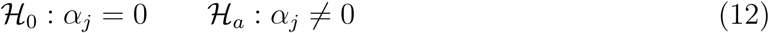

A likelihood ratio test (LRT) statistic for the *j*-th gene is given by

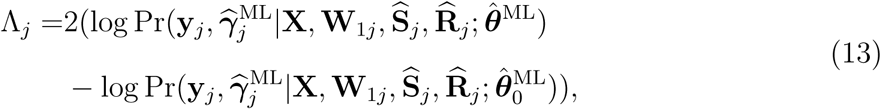

where 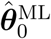 and 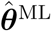 are vectors of parameter estimates that are obtained by maximizing the marginal likelihood, under the null hypothesis ℋ_0_ and under the alternative hypothesis ℋ_*A*_, respectively. Using standard asymptotic theory [38], the test statistics Λ_*j*_ asymptotically follows the 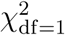 under the null.

As discussed in Section 2.3, to overcome the intractability of maximizing the marginal likelihood, we utilize a (PX)-VBEM algorithm where we maximize the ELBO, instead of the marginal likelihood, to obtain parameter estimates 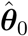 and 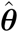. Earlier applications demonstrate that (PX)-VBEM produces practically useful and accurate posterior mean estimates [3, 8, 43, 34] (i.e. 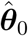 and 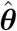). While it might seem reasonable to use the estimated posterior distribution from maximizing the ELBO to directly approximate the marginal likelihood in Equation 13, it is well-known that the (PX)-VBEM typically identifies posterior distributions that underestimate the marginal variances [40, 36]. Consequently, this is not a feasible approach. Instead of using the estimated posterior distribution as a proxy for the marginal likelihood in Equation 13, under the assumption that the posterior means of interest are well-estimated by (PX)-VBEM for all the parameters of interest (i.e. 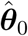 and 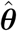 estimated using (PX)-VBEM are well-estimated), we plug-in our estimates from the (PX)-VBEM algorithm into the marginal likelihood in Equation 13 to construct the test statistic. We term the resulting likelihood with plug-in estimates from the (PX)-VBEM algorithm a calibrated ELBO and provide more details in the next section. Briefly, the calibrated ELBO is used as a proxy to the marginal likelihood in the test statistics. Our numerical studies show that a test constructed using the calibrated ELBO works well.

### 3.2 Calibrated ELBO

We postulate that the ELBO can be calibrated using the form from the (PX)-EM algorithm by plugging the posterior mean estimates and parameter estimates from (PX)-VBEM, which can be used as a proxy to marginal log-likelihood. Here we describe the procedures to calibrate the ELBO in detail. The marginal log-likelihood from the PX-EM algorithm for model (8) can be written as follows

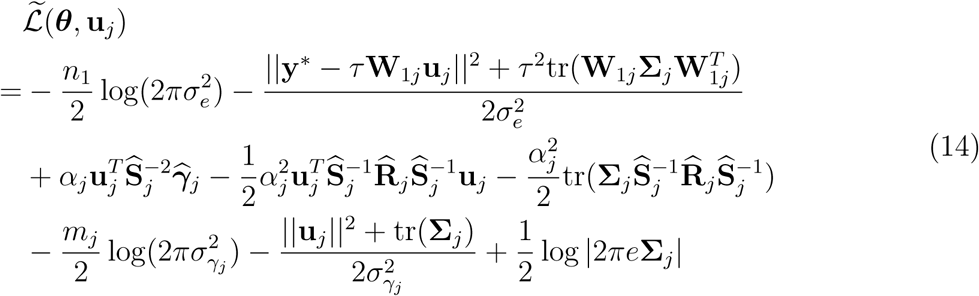

where **u**_*j*_ and **Σ**_*j*_ are the joint posterior mean and posterior variance for latent variable ***γ***_*j*_, and **Σ**_*j*_ is expressed as

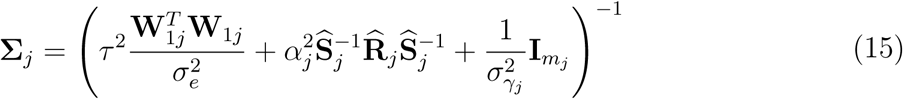

Note that we explicitly express the marginal log-likelihood (14) from the PX-EM algorithm that depends on the posterior mean, **u**_*j*_, and parameter estimates, ***θ***, as the posterior variance **Σ**_*j*_ is fully characterized by model parameters ***θ***. Thus, we first fit the data using the PX-VBEM (Algorithm 1 in the supplementary document). Then, we re-evaluate the marginal log-likelihood from the (PX)-EM algorithm by plugging the posterior mean estimates and parameter estimates 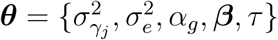 from the PX-VBEM algorithm as equation (14). Given the fact that the posterior means from (PX)-VBEM are accurate enough, the calibrated ELBO is close to the marginal log-likelihood. In Section 4, we conducted simulation studies to show that the marginal log-likelihood evaluated under the proposed calibrated procedure approximates well to that from (PX)-EM algorithms.

## 4. Simulations

### 4.1 Simulation settings

We conducted simulation studies to demonstrate that (a) CoMM-S^2^ has comparable performance as CoMM (Section 4.2.1) and that (b) CoMM-S^2^ generally performs as well or better than competing methods that also utilize summary statistics (Section 4.2.2). For (b), we compared the performance of CoMM-S^2^ with S-PrediXcan, with both ridge regression and Enet [47], denoted as S-PrediXcan:Ridge and S-PrediXcan:Enet, respectively.

We considered the following simulation settings to evaluate the performance of CoMM-S^2^. We assumed sample sizes of *n*_1_ = 400, *n*_2_ = 5, 000, and *n*_3_ = 400, which are the sample sizes for the transcriptome dataset, GWAS dataset and the reference panel dataset, respectively. To generate genotype data, we first generated a data matrix using a multivariate normal distribution 𝒩 (**0, Σ**(*ρ*)), where **Σ**(*ρ*) is an auto-regressive correlation structure with *ρ* = 0.2, 0.5 and 0.8, representing weak, moderate and strong LD, respectively. We then generated minor allele frequencies from a uniform distribution 𝒰 (0.05, 0.5) and categorized the data matrix into trinary variables taking values 0, 1, 2 using these minor allele frequencies and assuming Hardy-Weinberg equilibrium. All three genotype matrices, **W**_1*j*_, **W**_2*j*_ and 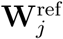 (where **W**_1*j*_, **W**_2*j*_ are as defined in Section 2.1, and 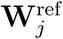 is a genotype data from a reference panel as described in Section 2.4) are generated in this manner.

To generate eQTL data from genotype data, we considered different cellular-level heritability levels 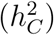 and sparsity levels, which are parameters that describe the genetic architecture of gene expression [41]. The cellular-level heritability 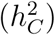 represents the proportion of variance of the eQTL that can be explained by genotype, while sparsity represents the proportion of genetic variants that are associated with the gene expression. For a given cellular-level heritability 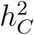, a larger number of genetic variants that are associated with gene expression levels implies a smaller genetic influence on gene expression, per genetic variant. We generated eQTL data assuming **y**_*j*_ = **W**_1*j*_***γ***_*j*_ + **e**_1_, where 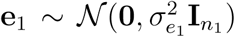, and the non-zero ***γ***_*j*_ were generated assuming 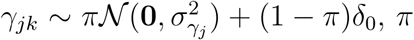 is the eQTL sparsity level, *δ*_0_ denotes a Dirac delta mass function at 0, and *k* is the index for genetic variants within gene *j*. 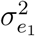 and 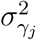 were chosen to correspond to cellular-level heritability levels 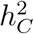 of 0.01, 0.05, or 0.09, which are close to the median gene expression heritability estimates computed across all genes [29]. We considered different eQTL sparsity levels of 0.1, 0.2, 0.3, 0.4 and 0.5, where a sparsity level of 0.2 indicates that only 20% of the SNPs have non zero effects (i.e. 20% of the *γ*_*j*_’s are non-zero).

We generated a complex trait assuming **z** = *α*_*j*_**W**_2*j*_***γ***_*j*_ + **e**_2_. We assumed 100 local genetic variants (cis-SNPs). **e**_2_ was chosen such that the organismal-level heritability level, defined as 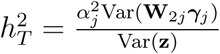 is controlled at 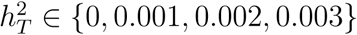. A organismal-level heritability 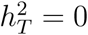 corresponds to the null hypothesis that the gene has no association with the organismal-level trait.

Summary statistics were generated by applying a single-variant analysis to the GWAS dataset.

### 4.2 Simulation results

#### 4.2.1 CoMM-S^2^ and CoMM have comparable performance

We first compared the LRT test statistics from both CoMM-S^2^ and ComMM. We used 1,000 simulation replicates to compare the LRT test statistics of the two methods As shown in Figures 1 and S2 - S7, the LRT test statistics of CoMM and CoMM-S^2^ are close to each other with a *R*^2^ around 0.98.

**Figure 1:**
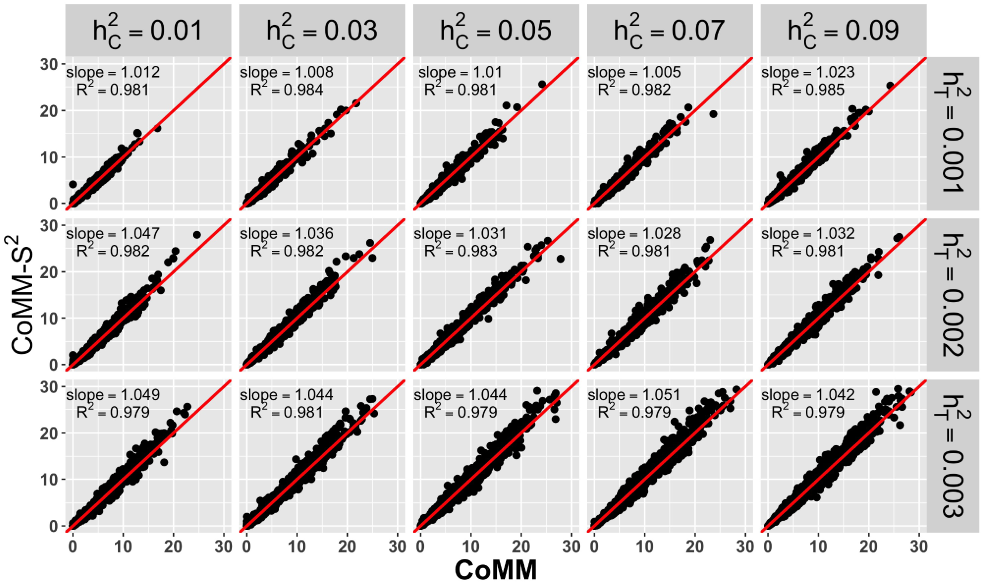
The scatter plot of test statistics from LRT for CoMM-S^2^ vs CoMM with the setting *n*_1_ = 400, *n*_2_ = 5,000, *n*_3_ = 400, *m*_*j*_ = 100, *ρ* = 0.8, *π* = 1. The number of replication is 2000. The reference panel is subsampled from GWAS dataset.

Next, we compared the calibrated ELBO as described in Section 3.2 with the marginal log-likelihood evaluated the using EM algorithm. Here, we consider the reference panel to be the GWAS data itself, which enables the evaluation of the marginal log-likelihood from EM algorithm. As shown in Figure S1 demonstrates that the calibrated ELBO is very similar to the marginal log-likelihood.

In the real data analysis of the NFBC1966 dataset (Section 5.2), we observed that a small proportion of test statistics from CoMM are degenerate zero. To better understand this phenomenon, we conducted additional simulations described in Supplementary Section 4.3. As shown in Figures S8 (a) and (b), while test statistics from CoMM degenerate to zero when 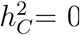, CoMM-S^2^ performs adequately under this setting.

#### 4.2.2 CoMM-S^2^ generally has better or comparable performance compared with alternative methods that use summary statistics

We evaluated the performance of CoMM-S^2^, S-PrediXcan:Ridge and S-PrediXcan:Enet under the null hypothesis 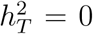, using 1,000 simulation replicates. The corresponding qq-plots are shown in Figures 2 and S9 - S11. The results show that CoMM-S^2^ can effectively control the type-I error, while S-PrediXcan shows a deflation when cellular heritability is low (*e.g.*, 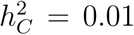). We also compared the power of the three methods. As shown in Figures 3 and S12, the power of all three methods increases as the cellular heritability 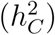 increases. CoMM-S^2^ generally outperforms S-PrediXcan (both Enet and Ridge) at low or moderate levels of cellular heritability (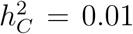 or 0.05). CoMM-S^2^ and S-PrediXcan:Ridge have comparable performance for larger values of 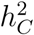. S-PrediXcan:Enet generally performs well at high levels of cellular heritability and when sparsity is low. The results also indicate that while the performance of CoMM-S^2^ and S-PrediXcan:Ridge do not vary very much with the sparsity, the performance of S-PrediXcan:Enet depends on the sparsity levels.

**Figure 2:**
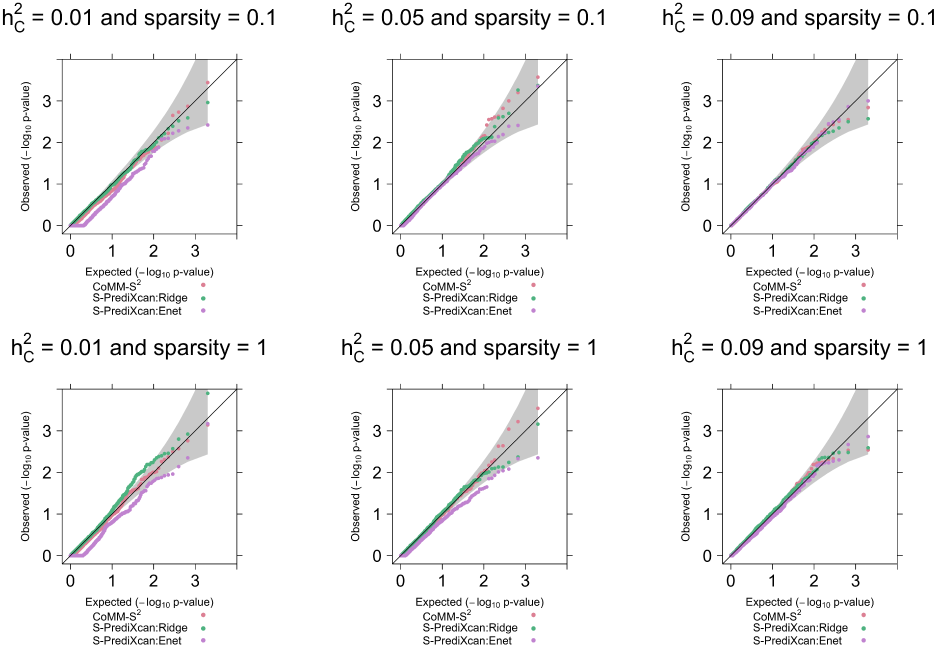
The qq-plot of *p*-values for each method (CoMM, PrediXcan:Ridge, PrediX-can:Enet, CoMM-S^2^, S-PrediXcan:Ridge and S-PrediXcan:Enet) with the setting *n*_1_ = 400, *n*_2_ = 5,000, *n*_3_ = 400, *ρ* = 0.8. The number of replication is 1000. The sparsity varies *π* ∈ {0.1, 1} and the cellular-level heritability varies from 0.01, 0.05, to 0.09.

**Figure 3:**
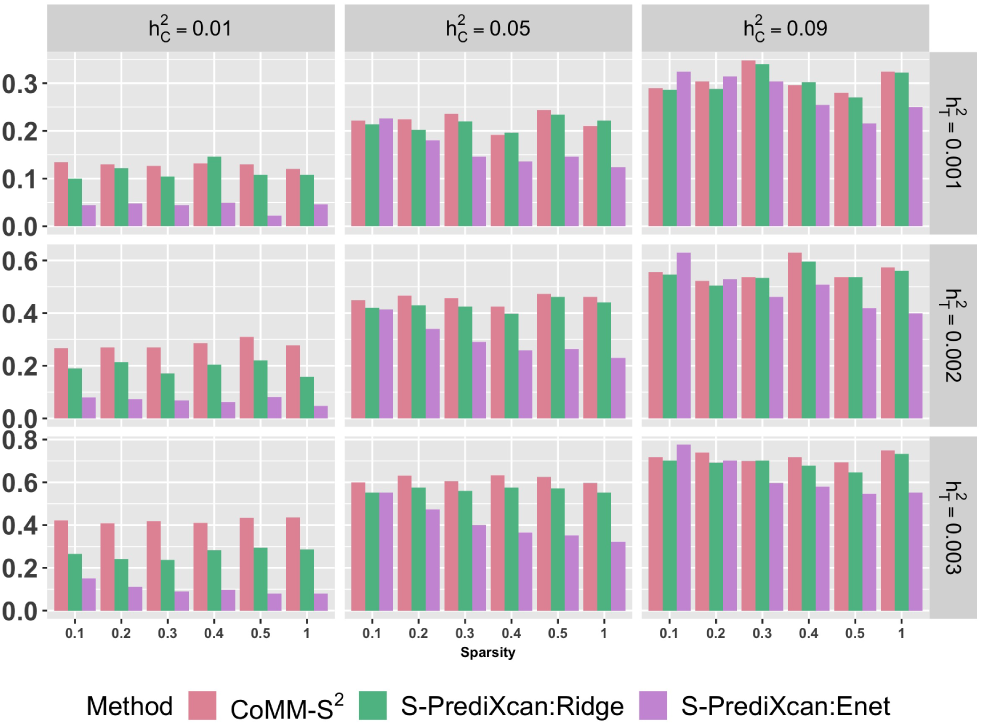
The comparison of power for CoMM, CoMM-S^2^, PrediXcan:Ridge, PrediX-can:Enet, S-PrediXcan:Ridge and S-PrediXcan:Enet with the setting *n*_1_ = 400, *n*_2_ = 5,000, *n*_3_ = 400, *ρ* = 0.8. The number of replication is 500. For each subplot, the x-axis stands for the sparsity of SNP and the y-axis stands for the proportion of significant genes within 500 replications.

## 5 Real Data Analysis

We applied CoMM-S^2^ to two data sets, individual-level data NFBC1966 [30] and summary statistics from 14 traits (see Tables S1 and S2), with the transcriptome data from GEU-VADIS Project [21] and Genotype-Tissue Expression (GTEx) Project [23], respectively. The NFBC dataset consists of information on ten quantitative traits. The ten quantitative traits include body mass index (BMI), systolic blood pressure (SysBP), diastolic blood pressure (DiaBP), high-density lipoprotein cholesterol (HDL-C), low-density lipoprotein cholesterol (LDL-C), triglycerides (TG), total cholesterol (TC), insulin levels, glucose levels and C-reactive protein (CRP). We also collected fourteen traits/diseases from multiple GWAS consortia including six diseases traits and eight BMI-related traits. Diseases traits are Alzheimer’s disease (AD), coronary artery disease (CAD), autism spectrum disorder (ASD), schizophrenia (SCZ), type 2 diabetes (T2D), myocardial infarction (MI). There are eight summary statistics for BMI-related traits from 2017 GIANT Gene-Physical Activity Interaction Meta-analysis [12] including two sex-combined traits, BMI for physically active individuals (BMIPA), BMI for physically inactive individuals (BMIPI), and other six sex-specific traits, BMI for physically active individuals in men(BMIPAM), BMI for physically active individuals in women(BMIPAW), BMI for physically inactive individuals in men(BMIPIM), BMI for physically inactive individuals in women(BMIPIW), BMI adjusted for physical activity for individuals in men(BMIadjPAM) and BMI adjusted for physical activity for individuals in women(BMIadjPAW). For transcriptome data, GEUVADIS study contains 15,810 genes and GTEx project has gene expressions for 48 tissues, where the number of genes in each tissue ranges from 16,333 to 27,378.

### 5.1 Analysis of NFBC1966 datatset

As we have individual-level data for NFBC1966 dataset, we then applied both CoMM and CoMM-S^2^ using individual-level data and summary statistics, respectively. We first analyzed the individual-level data from NFBC1966 with the transcriptome data from GEUVADIS using CoMM. The results from CoMM can be taken as a benchmark as it uses individual-level data. We then conducted single-variant analysis for NFBC1966 dataset to generate summary statistics. Finally, we applied CoMM-S^2^ using summary statistics from NFBC1966 dataset. As CoMM-S^2^ uses a reference panel to give estimates for LD, 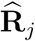, we applied two different choices of reference panel, namely, 400 subsamples from NFBC1966 and European samples from 1KG [7]. The scatter plots of LRT test statistics for CoMM against CoMM-S^2^ using 400 subsamples from NFBC1966 are given in Figure 4 and the one using 1KG samples as reference data is shown in Figure S24 in the supplementary document. When we used the subsamples as the reference panel (in Figure 4), we notice that the test statistics from CoMM-S^2^ are close to their counterpart with slope around 1 and *R*^2^ ranging from 0.91 to 0.99. When the reference panel becomes 1KG as shown in Figure S13, one can observe that the test statistics in the null region (Λ_*g*_ < 20.84 *≡ p*-value ¿ 5 × 10^−6^) from both CoMM and CoMM-S^2^ are roughly around the line with slope equals to 1. The test statistics in the non-null region (Λ_*g*_ ≥ 20.84 *≡ p*-value ≤ 5 × 10^−6^) are inflated. This is primarily due to the reference panel we used to estimate correlations. When we applied sub-samples from NFBC1966 as reference panel data, this difference essentially disappeared. The reason for this phenomenon could be that despite that NFBC1966 dataset is a Finn’s study from Europe, Finnish samples was shown its genetic distinctness in previous studies [31].Although this genetic discrepancy for Finnish population, we found that the inflation only appears in the non-null regions, which makes the use of 1KG as reference panel practically useful. Note that in these comparisons, we removed genes with cellular heritability less than 0.01 as the test statistics for tiny cellular heritability using CoMM is not reliable. The use of cutoff here is to make fair comparisons between CoMM and CoMM-S^2^ as test statistics of CoMM degenerates to zero when cellular heritability is small (see Figure S8). In practice, we do not require this limit as CoMM-S^2^ also works in tiny cellular heritability regions. To verify the phenomenon of degeneration, we compared CoMM-S^2^ and CoMM with RL-SKAT [33] for genes with cellular heritability less than 0.01 as shown in Figure S14. The result is consistent with the simulation results from both Section 4.2 and our previous work [42].

**Figure 4:**
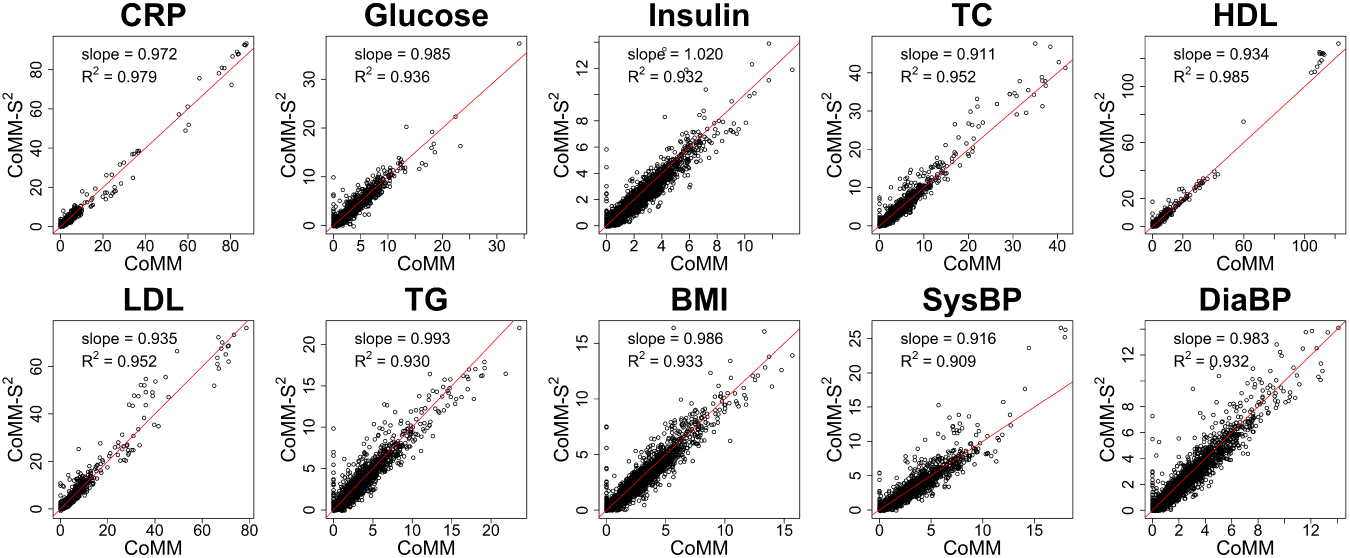
The scatter plot of LRT statistics for CoMM-IS vs CoMM, with transcriptiome data from GTEx tissue Adipose Subcutaneous, GWAS data from NFBC1966 dataset, and reference panel data from 400 subsamples of NFBC1966 dataset. We remove the points with cellular heritability less than 0.001.

### 5.2 Analysis of 14 traits

We performed the analysis for 14 GWAS summary statistics with transcriptome data from GTEx and the detailed information of these 14 traits are shown in Tables S1 and S2. Specifically, we applied CoMM-S^2^ together with S-PrediXcan (both Enet and Ridge) to examine the associations between each pair of a gene and the complex trait. We display qq-plots for each trait across all tissues in Figure S15. After completing the analysis using three approaches, we conducted genomic control for each trait-tissue pair. The genome-wide significant threshold is set to be 5 × 10^−6^ based on Bonferroni correction.

The results indicate that CoMM-S^2^ identified more significant associations than S-PrediXcan. The analysis for each individual summary statistics together with the transcriptome data for a tissue can be done around 20 min on a Linux platform with 2.6 GHz intel Xeon CPU E5-2690 with 30 720 KB cache and 96 GB RAM (0nly 6∼7 GB RAM used) on 24 cores.

CoMM-S^2^ is primarily developed to identify the associations between genes and complex traits using summary statistics. It is not only power in the region that cellular heritability is relatively large, but can also identify associations in the weak cellular heritability region. We show both the number of unique genes passing genome-significance level across different tissues and the number of genes reported in the previous studies in Table 1 while the total number of genes identified across 48 tissues using CoMM-S^2^ and S-PrediXcan (both Enet and Ridge) is shown in Table S3. For example, as shown in Table 1, CoMM-S^2^ identified 199 unique genes across all tissues and 37 of them were previously reported in NHGRI-EBI GWAS Catalog [4] for AD while S-PrediXcan (both Enet and Ridge) identified 95 and 108 genes with 24 and 28 genes reported before, respectively. However, CoMM-S^2^ identified 4,614 genes in total across 48 tissues while S-PrediXcan (both Enet and Ridge) identified only 398 and 664 genes in total, respectively, which indicates that there are more overlapped genes identified by CoMM-S^2^ but the genes identified by S-PrediXcan (both Enet and Ridge) are more or less unique across tissues. Specifically, in GTEx adipose subcutaneous tissue, CoMM-S^2^ identified seven genes in band 2q14.3, 12 genes in band 8p21.2, 14 genes in bands 11q12.1 and 11q12.2, 13 genes in bands 11q14.1 and 11q14.2, and 51 genes in band 19q13.32. Among them, 24 genes were reported to be associated with AD in previous studies [5, 6, 24, 9, 28, 19]. However, S-PrediXcan can only identify six of these 24 genes. Note that 18 out of these 24 genes have 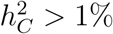 and among the six genes having 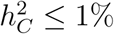, S-PrediXcan (both Enet and Ridge) only identified one gene and none, respectively. In addition, gene *BIN1* was identified among all 48 tissues and cellular heritability for gene *BIN1* are larger than 10% in eight tissues, *e.g.*, 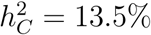 in brain cerebellum, 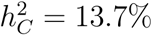 in brain cortex, 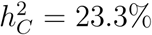 in esophagus muscularis mucosa, and highest in pancreas with 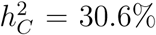. Hence, the associations between gene *BIN1* and AD in tissues with large 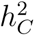 are more likely to be causal, where targeting *BIN1* might present novel AD therapy [35]. Gene *PTK2B* has largest cellular heritability in tissue brain cerebellum 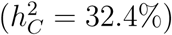, which was one of most significant genes associated with AD in a meta-analysis [20]. In a mouse model, overexpression of *PTK2B* improved the behavioral and molecular phenotype of a strain of AD-linked mutated mice [11]. Gene *CLU* has the largest cellular heritability in the adrenal gland tissue 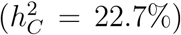, which has been found to be related to cholesterol synthesis, transport, uptake or metabolism in AD that links between cholesterol and AD pathogenesis [15].

**Table 1:** The number of significant genes identified across the tissues at the significant level (5 ×10^−6^) for each method. In this table, the same genes identified across the tissues are only counted once. The number within the parenthesis denoted the number of genes reported in NHGRI-EBI GWAS Catalog [4].

|  | CoMM-S<sup>2</sup> | S-PrediXcan:Enet | S-PrediXcan:Ridge |
| --- | --- | --- | --- |
| AD | 199(37) | 105(28) | 111(29) |
| ASD | 2(0) | 1(0) | 5(0) |
| CAD | 117(15) | 71(11) | 73(11) |
| MI | 119(6) | 81(7) | 51(7) |
| SCZ | 61(11) | 57(17) | 53(10) |
| T2D | 160(27) | 109(27) | 110(22) |
| BMIPA | 258(54) | 137(35) | 126(36) |
| BMIPI | 53(11) | 22(7) | 28(7) |
| BMIPAM | 31(4) | 9(1) | 17(4) |
| BMIPAW | 123(31) | 68(18) | 67(21) |
| BMIPIM | 12(2) | 8(2) | 7(4) |
| BMIPIW | 24(4) | 13(3) | 11(2) |
| BMIadjPAM | 104(21) | 54(16) | 76(21) |
| BMIadjPAW | 208(45) | 102(31) | 107(30) |

For CAD, CoMM-S^2^ identified 117 unique genes across all tissues and 15 of them were previously reported in NHGRI-EBI GWAS Catalog [4] while S-PrediXcan (both Enet and Ridge) identified 71 and 73 genes with 11 and 11 genes reported before, respectively. However, CoMM-S^2^ identified 2,067 genes in total across all tissues while S-PrediXcan (both Enet and Ridge) identified only 109 and 184 genes in total, respectively. Specifically, in the artery aorta tissue, CoMM-S^2^ identified two genes in band 2q33.2, one gene in band 3q22.3, three genes in band 6p24.1, two genes in band 6q25.3, eight genes in band 9p21.3, four genes in bands 12q24.11 and 12q24.12, six genes in band 13q34, seven genes in band 15q25.1, and five genes in band 19p13.2. Among them, nine genes were reported to be associated with CAD in previous studies [14, 25, 37]. However, S-PrediXcan can only identify two of these nine genes. Gene *NBEAL1* was identified to be genome-wide significant in eleven tissues. Among these eleven tissues, 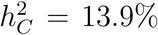 in artery aorta, 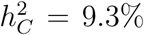 in artery tibial, and 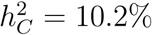 in pituitary are relative large, in which gene *NBEAL1* is likely to be causal for CAD.

The results of the identified genes for all 14 traits across tissues together with their corresponding test statistics and *p*-values for CoMM-S^2^, S-PrediXcan (both Enet and Ridge) can be found in excel tables in the supplementary files.

## 6 Conclusion

In this article, we have developed a collaborative mixed model using summary statistics from GWAS to account for uncertainty in transcriptome imputation. We examined the relationship between CoMM and CoMM-S^2^. Our numerical results show that CoMM-S^2^ has comparable performance as CoMM. CoMM-S^2^ has several advantages over CoMM. First, CoMM-S^2^ can be computationally more efficient than CoMM when applied to GWAS that have large sample sizes. This is because CoMM-S^2^ is applied to summary statistics, which is faster to compute in large sample sizes, while CoMM is applied to individual-level data. Second, through empirical studies, we show that CoMM-S^2^ has better performance when the cellular heritability is low. However, CoMM-S^2^ is not without limitations. First, CoMM-S^2^ cannot be utilized in a cross-tissue analysis, for example as illustrated in [17]. Furthermore, CoMM-S^2^ cannot differentiate whether the identified genes are simply associated with the complex traits or if they are real causal effects. These are avenues for further research.

