## supplementary for "CoMM-S^2^: a collaborative mixed model using summary statistics in transcriptome-wide association studies" (from another version of this preprint: version 1, 2019-05-29)

### 1 Data Sources

#### 1.1 GWAS data

We analyzed two sets of data sets, individual-level data NFBC1966 [12] and summary statistics from 14 traits (see Tables S1 and S2). The NFBC1966 data set contains information for 5,402 individuals with a selected list of phenotypic data related to cardiovascular disease including multiple metabolic traits (high-density lipoprotein cholesterol, HDL-C; low-density lipoprotein cholesterol, LDL-C; total cholesterol, TC; triglycerides, TG; C-reactive protein, CRP), glucose, insulin, body mass index (BMI), systolic and diastolic blood pressure (SysBP and DiaBP), and 364,590 SNPs genotyped.

We performed strict quality control on data using PLINK [11]. We excluded individuals having discrepancies between reported sex and sex determined from the X chromosome. We excluded SNPs with a minor allele frequency less than 1%, having missing values in more than 1% of the individuals or with a Hardy-Weinberg equilibrium p-value below 0.0001. After quality control, 5,123 individuals with 319,147 SNPs in NFBC1966.

#### 1.2 Transcriptome data

The Genetic European Variation in Health and Disease (GEUVADIS) Project [8] provided mRNA and small RNA sequencing on 465 Epstein-Barr-virus-transformed lymphoblastoid cell lines samples whose DNA sequencing data can be available from the 1000 Genomes Project [5]. In this study, we only used 379 individuals from European ancestry in the 1000 Genome Project, including British in England and Scotland, Finnish in Finland, Toscani in Italy and Utah residents with Northern and Western European ancestry. Since the original gene expression measurements are read counts , the PEER normalization [16, 17, 1] was applied to remove technical variations and batch effects. We quantile-normalized every gene expression to a standard normal distribution separately in the five populations and then quantile-normalized them together. According to GENCODE (release 12) [7], we selected 15,810 genes in the GEUVADIS data set that were expressed in at least half of the individuals. For each gene, we only included the common (minor allele frequency >= 0.05) variants within 50kb of the transcription start and end of each protein coding gene (according to GENCODE release 12 gene annotation), resulting in 550 variants on average included for one gene.

The Genotype-Tissue Expression (GTEx) Project [9] began to provide DNA sequencing data (about 12.5 million variants) from 449 individuals and collected gene-expression measurements of 44 tissues from these individuals. Now GTEx contains 714 samples from 53 tissues in release V7, among them 48 tissues have eQTL data. Data repository for GTEx (dbGaP Study Accession: phs000424.v7.p2) has already performed gene normalization for RNA-seq data and provides user with Transcripts Per Million (TPM) data. In total, GTEx contains gene expressions for 48 tissues and the number of genes within a tissue ranges from 16,333 to 27,378. For the genotype data in GTEx, we conducted quality control to include the common (minor allele frequency >= 0.05) variants within 50kb of the transcription start and end of each protein coding gene, resulting in 400 variants on average included for one gene.

### 2 PX-VBEM Algorithm

In this section, we provide technical details for the algorithm to solve the model detailed in Section 2 of main text. In Section 2.1, we first show the derivation for the variational posterior distribution of latent variables ($\boldsymbol{\gamma}_j$) following by the derivation of ELBO of the model (4) by variational expectations in Section 2.2. Then, in Section 2.3, we show the maximum likelihood estimates (MLE) of the model parameters ($\boldsymbol{\theta}$) by setting the first order derivative of ELBO to be 0. Finally, we show the overall PX-VBEM algorithm in Section 2.4.

#### 2.1 Variational posterior distribution

The logarithm of complete-data likelihood (4) can be written as

$$
\begin{aligned}
&\Pr(\mathbf{y}_j, \widehat{\boldsymbol{\gamma}}_j, \boldsymbol{\gamma}_j | \mathbf{X}, \mathbf{W}_{1j}, \widehat{\mathbf{S}}_j, \widehat{\mathbf{R}}_j; \boldsymbol{\theta}) \\
=&- \frac{n_1}{2}\log(2\pi\sigma_e^2) - \frac{||\mathbf{y}_j^* - \tau\mathbf{W}_{1j}\boldsymbol{\gamma}_j||^2}{2\sigma_e^2} \\
&- \frac{1}{2}\log|\widehat{\mathbf{S}}_j\widehat{\mathbf{R}}_j\widehat{\mathbf{S}}_j| - \frac{1}{2}(\widehat{\boldsymbol{\gamma}}_j - \alpha_j\widehat{\mathbf{S}}_j\widehat{\mathbf{R}}_j\widehat{\mathbf{S}}_j^{-1}\boldsymbol{\gamma}_j)^\top(\widehat{\mathbf{S}}_j\widehat{\mathbf{R}}_j\widehat{\mathbf{S}}_j)^{-1}(\widehat{\boldsymbol{\gamma}}_j - \alpha_j\widehat{\mathbf{S}}_j\widehat{\mathbf{R}}_j\widehat{\mathbf{S}}_j^{-1}\boldsymbol{\gamma}_j) \\
&- \frac{m_j}{2}\log(2\pi\sigma_{\gamma_j}^2) - \frac{||\boldsymbol{\gamma}_j||^2}{2\sigma_{\gamma_j}^2},
\end{aligned}
\tag{S1}
$$

where $\mathbf{y}_j^* = \mathbf{y}_j - \mathbf{X}\boldsymbol{\beta}$. The terms involving $\gamma_{jk}$ in the log-complete-data likelihood (S1) has the following quadratic form,

$$
\begin{aligned}
&-\frac{1}{2}\left(\frac{\tau^2}{\sigma_e^2}\mathbf{w}_{1jk}^\top\mathbf{w}_{1jk} + \alpha_j^2\frac{R_{jkk}}{S_{jkk}^2} + \frac{1}{\sigma_{\gamma_j}^2}\right)\gamma_{jk}^2 \\
&+\left(\frac{\tau}{\sigma_e^2}(\mathbf{y}_j^* - \tau\sum_{i\neq k}\gamma_{ji}\mathbf{w}_{1ji})^\top\mathbf{w}_{1jk} + \alpha_j\frac{\widehat{\gamma}_{jk}}{S_{jkk}^2} - \alpha_j^2\sum_{i\neq k}\frac{\gamma_{ji}R_{jki}}{S_{jii}S_{jkk}}\right)\gamma_{jk}.
\end{aligned}
\tag{S2}
$$

Referring to the formula (10.9) in Bishop's book [3], taking the expectation under the distribution $q(\boldsymbol{\gamma}_{ji}), i \neq k$ and the expression of $\log q(\gamma_{jk})$ can be derived as follows

$$
\begin{aligned}
\log q(\gamma_{jk}) =& -\frac{1}{2}\left(\frac{\tau^2}{\sigma_e^2}\mathbf{w}_{1jk}^\top\mathbf{w}_{1jk} + \alpha_j^2\frac{R_{jkk}}{S_{jkk}^2} + \frac{1}{\sigma_{\gamma_j}^2}\right)\gamma_{jk}^2 \\
&+\left(\frac{\gamma_j}{\sigma_e^2}(\mathbf{y}_j^* - \tau\sum_{i\neq k}\mathbb{E}[\gamma_{ji}]\mathbf{w}_{1ji})^\top\mathbf{w}_{1jk} + \alpha_j\frac{\widehat{\gamma}_{jk}}{S_{jkk}^2} - \alpha_j^2\sum_{i\neq k}\frac{\mathbb{E}[\gamma_{ji}]R_{jki}}{S_{jii}S_{jkk}}\right)\gamma_{jk}.
\end{aligned}
\tag{S3}
$$

Clearly, the posterior of $\gamma_{jk}$ is a Gaussian distribution $\mathcal{N}(u_{jk}, v_{jk}^2)$, where the mean $u_{jk}$ and variance $v_{jk}^2$ are defined as

$$
\begin{aligned}
v_{jk}^2 &= \frac{1}{\frac{\tau^2}{\sigma_e^2}\mathbf{w}_{1jk}^\top\mathbf{w}_{1jk} + \alpha_j^2\frac{R_{jkk}}{S_{jkk}^2} + \frac{1}{\sigma_{\gamma_j}^2}}, \\
u_{jk} &= \frac{\frac{\gamma_j}{\sigma_e^2}(\mathbf{y}_j^* - \tau\sum_{i\neq k}\mathbb{E}[\gamma_{ji}]\mathbf{w}_{1ji})^\top\mathbf{w}_{1j} + \alpha_j\frac{\widehat{\gamma}_{jk}}{S_{jkk}^2} - \alpha_j^2\sum_{i\neq k}\frac{\mathbb{E}[\gamma_{ji}]R_{jki}}{S_{jii}S_{jkk}}}{\frac{\tau^2}{\sigma_e^2}\mathbf{w}_{1jk}^\top\mathbf{w}_{1jk} + \alpha_j^2\frac{R_{jkk}}{S_{jkk}^2} + \frac{1}{\sigma_{\gamma_j}^2}}.
\end{aligned}
\tag{S4}
$$

#### 2.2 ELBO and VB E-step

Now we can evaluate the ELBO $\mathcal{L}(q)$, it is defined as

$$\mathcal{L}(q) = \mathbb{E}_q[\log \Pr(\mathbf{y}_j, \widehat{\boldsymbol{\gamma}}_j, \boldsymbol{\gamma}_j | \mathbf{X}, \mathbf{W}_{1j}, \widehat{\mathbf{S}}_j, \widehat{\mathbf{R}}_j; \boldsymbol{\theta})] - \mathbb{E}_q[\log q(\boldsymbol{\gamma}_j)]. \tag{S5}$$

The first term of the ELBO is the expectation of log-complete-data likelihood evaluated under the variational distribution, which can be written as

$$\begin{aligned} &\mathbb{E}_q[\log \Pr(\mathbf{y}_j, \widehat{\boldsymbol{\gamma}}_j, \boldsymbol{\gamma}_j | \mathbf{X}, \mathbf{W}_{1j}, \widehat{\mathbf{S}}_j, \widehat{\mathbf{R}}_j; \boldsymbol{\theta})] \\ =& -\frac{n_1}{2}\log(2\pi\sigma_e^2) - \frac{\mathbb{E}_q||\mathbf{y}_j^* - \tau\mathbf{W}_{1j}\boldsymbol{\gamma}_j||^2}{2\sigma_e^2} \\ &- \frac{1}{2}\log|\widehat{\mathbf{S}}_j\widehat{\mathbf{R}}_j\widehat{\mathbf{S}}_j| - \frac{1}{2}\mathbb{E}_q[(\widehat{\boldsymbol{\gamma}}_j - \alpha_j\widehat{\mathbf{S}}_j\widehat{\mathbf{R}}_j\widehat{\mathbf{S}}_j^{-1}\boldsymbol{\gamma}_j)^\top(\widehat{\mathbf{S}}_j\widehat{\mathbf{R}}_j\widehat{\mathbf{S}}_j)^{-1}(\widehat{\boldsymbol{\gamma}}_j - \alpha_j\widehat{\mathbf{S}}_j\widehat{\mathbf{R}}_j\widehat{\mathbf{S}}_j^{-1}\boldsymbol{\gamma}_j)] \\ &- \frac{m_j}{2}\log(2\pi\sigma_{\gamma_j}^2) - \frac{\mathbb{E}_q||\boldsymbol{\gamma}_j||^2}{2\sigma_{\gamma_j}^2}. \end{aligned} \tag{S6}$$

In this expression, it involves the quantities for related moments under variational distribution, which can be written as

$$\begin{aligned} &\mathbb{E}_q[\boldsymbol{\gamma}_j] = \mathbf{u}_j, \\ &\mathbb{E}_q\|\boldsymbol{\gamma}_j\|^2 = \mathbf{u}_j^\top\mathbf{u}_j + \mathbf{v}_j^\top\mathbf{v}_j, \\ &\mathbb{E}_q[\boldsymbol{\gamma}_j^\top\mathbf{W}_{1j}^\top\mathbf{W}_{1j}\boldsymbol{\gamma}_j] = \mathbf{u}_j^\top\mathbf{W}_{1j}^\top\mathbf{W}_{1j}\mathbf{u}_j + \sum_{k=1}^{m_j}\mathbf{w}_{1jk}^\top\mathbf{w}_{1jk}v_{jk}^2, \\ &\mathbb{E}_q[\boldsymbol{\gamma}_j^\top\widehat{\mathbf{S}}_j^{-1}\widehat{\mathbf{R}}_j\widehat{\mathbf{S}}_j^{-1}\boldsymbol{\gamma}_j] = \mathbf{u}_j^\top\widehat{\mathbf{S}}_j^{-1}\widehat{\mathbf{R}}_j\widehat{\mathbf{S}}_j^{-1}\mathbf{u}_j + (\mathbf{v}_j \odot \mathbf{v}_j)^\top\mathrm{diag}(\widehat{\mathbf{S}}_j^{-1}\widehat{\mathbf{R}}_j\widehat{\mathbf{S}}_j^{-1}), \end{aligned} \tag{S7}$$

where the notation $\odot$ denotes the element-wise multiplication of two vectors and diag($\mathbf{A}$) denotes a vector whose elements are extracted from the diagonal entries of the matrix $\mathbf{A}$. Using these quantities, we can evaluate the two terms in expression (S6) as

$$\begin{aligned} \mathbb{E}_q||\mathbf{y}_j^* - \tau\mathbf{W}_{1j}\boldsymbol{\gamma}_j||^2 =& \mathbb{E}_q[\mathbf{y}_j^{*\top}\mathbf{y}_j^* - 2\tau\mathbf{y}_j^{*\top}\mathbf{W}_{1j}\boldsymbol{\gamma}_j + \tau^2\boldsymbol{\gamma}_j^\top\mathbf{W}_{1j}^\top\mathbf{W}_{1j}\boldsymbol{\gamma}_j] \\ =& \mathbf{y}_j^{*\top}\mathbf{y}_j^* - 2\tau\mathbf{y}_j^{*\top}\mathbf{W}_{1j}\mathbf{u}_j + \tau^2\mathbf{u}_j^\top\mathbf{W}_{1j}^\top\mathbf{W}_{1j}\mathbf{u}_j + \tau^2\sum_{k=1}^{m_j}\mathbf{w}_{1jk}^\top\mathbf{w}_{1jk}v_{jk}^2 \\ =& ||\mathbf{y}_j^* - \tau\mathbf{W}_{1j}\mathbf{u}_j||^2 + \tau^2\sum_{k=1}^{m_j}\mathbf{w}_{1jk}^\top\mathbf{w}_{1jk}v_{jk}^2, \end{aligned} \tag{S8}$$

$$
\begin{aligned}
&\mathbb{E}_q[(\widehat{\boldsymbol{\gamma}}_j - \alpha_j \widehat{\mathbf{S}}_j \widehat{\mathbf{R}}_j \widehat{\mathbf{S}}_j^{-1} \boldsymbol{\gamma}_j)^\top (\widehat{\mathbf{S}}_j \widehat{\mathbf{R}}_j \widehat{\mathbf{S}}_j)^{-1} (\widehat{\boldsymbol{\gamma}}_j - \alpha_j \widehat{\mathbf{S}}_j \widehat{\mathbf{R}}_j \widehat{\mathbf{S}}_j^{-1} \boldsymbol{\gamma}_j)] \\
=&\mathbb{E}_q[\widehat{\boldsymbol{\gamma}}_j^\top (\widehat{\mathbf{S}}_j \widehat{\mathbf{R}}_j \widehat{\mathbf{S}}_j)^{-1} \widehat{\boldsymbol{\gamma}}_j - 2\alpha_j \boldsymbol{\gamma}_j^\top \widehat{\mathbf{S}}_j^{-2} \widehat{\boldsymbol{\gamma}}_j + \alpha_j^2 \boldsymbol{\gamma}_j^\top \widehat{\mathbf{S}}_j^{-1} \widehat{\mathbf{R}}_j \widehat{\mathbf{S}}_j^{-1} \boldsymbol{\gamma}_j] \\
=&\widehat{\boldsymbol{\gamma}}_j^\top (\widehat{\mathbf{S}}_j \widehat{\mathbf{R}}_j \widehat{\mathbf{S}}_j)^{-1} \widehat{\boldsymbol{\gamma}}_j - 2\alpha_j \mathbf{u}_j^\top \widehat{\mathbf{S}}_j^{-2} \widehat{\boldsymbol{\gamma}}_j + \alpha_j^2 \mathbf{u}_j^\top \widehat{\mathbf{S}}_j^{-1} \widehat{\mathbf{R}}_j \widehat{\mathbf{S}}_j^{-1} \mathbf{u}_j + \alpha_j^2 (\mathbf{v}_j \odot \mathbf{v}_j)^\top \mathrm{diag}(\widehat{\mathbf{S}}_j^{-1} \widehat{\mathbf{R}}_j \widehat{\mathbf{S}}_j^{-1}) \\
=&(\widehat{\boldsymbol{\gamma}}_j - \alpha_j \widehat{\mathbf{S}}_j \widehat{\mathbf{R}}_j \widehat{\mathbf{S}}_j^{-1} \mathbf{u}_j)^\top (\widehat{\mathbf{S}}_j \widehat{\mathbf{R}}_j \widehat{\mathbf{S}}_j)^{-1} (\widehat{\boldsymbol{\gamma}}_j - \alpha_j \widehat{\mathbf{S}}_j \widehat{\mathbf{R}}_j \widehat{\mathbf{S}}_j^{-1} \mathbf{u}_j) + \alpha_j^2 (\mathbf{v}_j \odot \mathbf{v}_j)^\top \mathrm{diag}(\widehat{\mathbf{S}}_j^{-1} \widehat{\mathbf{R}}_j \widehat{\mathbf{S}}_j^{-1}),
\end{aligned}
\tag{S9}
$$

The analytical form of ELBO can be obtained by plugging in these two terms,

$$
\begin{aligned}
&\mathbb{E}_q[\log \Pr(\mathbf{y}_j, \widehat{\boldsymbol{\gamma}}_j, \boldsymbol{\gamma}_j | \mathbf{X}, \mathbf{W}_{1j}, \widehat{\mathbf{S}}_j, \widehat{\mathbf{R}}_j; \boldsymbol{\theta})] \\
=& -\frac{n_1}{2} \log(2\pi\sigma_e^2) - \frac{||\mathbf{y}_j^* - \tau \mathbf{W}_{1j} \mathbf{u}_j||^2 + \tau^2 \sum_{k=1}^{m_j} \mathbf{w}_{1jk}^\top \mathbf{w}_{1jk} v_{jk}^2}{2\sigma_e^2} \\
&- \frac{1}{2} \log |\widehat{\mathbf{S}}_j \widehat{\mathbf{R}}_j \widehat{\mathbf{S}}_j| - \frac{1}{2} (\widehat{\boldsymbol{\gamma}}_j - \alpha_j \widehat{\mathbf{S}}_j \widehat{\mathbf{R}}_j \widehat{\mathbf{S}}_j^{-1} \mathbf{u}_j)^\top (\widehat{\mathbf{S}}_j \widehat{\mathbf{R}}_j \widehat{\mathbf{S}}_j)^{-1} (\widehat{\boldsymbol{\gamma}}_j - \alpha_j \widehat{\mathbf{S}}_j \widehat{\mathbf{R}}_j \widehat{\mathbf{S}}_j^{-1} \mathbf{u}_j) \\
&- \frac{\alpha_j^2}{2} (\mathbf{v}_j \odot \mathbf{v}_j)^\top \mathrm{diag}(\widehat{\mathbf{S}}_j^{-1} \widehat{\mathbf{R}}_j \widehat{\mathbf{S}}_j^{-1}) \\
&- \frac{m_j}{2} \log(2\pi\sigma_{\gamma_j}^2) - \frac{||\mathbf{u}_j||^2 + ||\mathbf{v}_j||^2}{2\sigma_{\gamma_j}^2}.
\end{aligned}
\tag{S10}
$$

So far, we still need evaluate the entropy term of ELBO,

$$
-\mathbb{E}_q[\log q(\boldsymbol{\gamma}_{jk})] = \frac{1}{2} \log |2\pi e v_{jk}^2| \tag{S11}
$$

Combing the two terms of ELBO (S5), we can obtain the following analytic form as

$$
\begin{aligned}
\mathcal{L}(q) =& -\frac{n_1}{2} \log(2\pi\sigma_e^2) - \frac{||\mathbf{y}_j^* - \tau \mathbf{W}_{1j} \mathbf{u}_j||^2 + \tau^2 \sum_{k=1}^{m_j} \mathbf{w}_{1jk}^\top \mathbf{w}_{1jk} v_{jk}^2}{2\sigma_e^2} \\
&- \frac{1}{2} \log |\widehat{\mathbf{S}}_j \widehat{\mathbf{R}}_j \widehat{\mathbf{S}}_j| - \frac{\alpha_j^2}{2} (\mathbf{v}_j \odot \mathbf{v}_j)^\top \mathrm{diag}(\widehat{\mathbf{S}}_j^{-1} \widehat{\mathbf{R}}_j \widehat{\mathbf{S}}_j^{-1}) \\
&- \frac{1}{2} (\widehat{\boldsymbol{\gamma}}_j - \alpha_j \widehat{\mathbf{S}}_j \widehat{\mathbf{R}}_j \widehat{\mathbf{S}}_j^{-1} \mathbf{u}_j)^\top (\widehat{\mathbf{S}}_j \widehat{\mathbf{R}}_j \widehat{\mathbf{S}}_j)^{-1} (\widehat{\boldsymbol{\gamma}}_j - \alpha_j \widehat{\mathbf{S}}_j \widehat{\mathbf{R}}_j \widehat{\mathbf{S}}_j^{-1} \mathbf{u}_j) \\
&- \frac{m_j}{2} \log(2\pi\sigma_{\gamma_j}^2) - \frac{||\mathbf{u}_j||^2 + ||\mathbf{v}_j||^2}{2\sigma_{\gamma_j}^2} + \frac{1}{2} \sum_{k=1}^{m_j} \log |2\pi e v_{jk}^2|.
\end{aligned}
\tag{S12}
$$

Note that the terms $-\frac{1}{2}\log|\widehat{\mathbf{S}}_j\widehat{\mathbf{R}}_j\widehat{\mathbf{S}}_j|$ and $\widehat{\boldsymbol{\gamma}}_j^\top(\widehat{\mathbf{S}}_j\widehat{\mathbf{R}}_j\widehat{\mathbf{S}}_j)^{-1}\widehat{\boldsymbol{\gamma}}_j$ are constant. Thus we can omit the constants in ELBO for simplicity,

$$
\begin{aligned}
\mathcal{L}(q) =& -\frac{n_1}{2}\log(2\pi\sigma_e^2) - \frac{||\mathbf{y}_j^* - \tau\mathbf{W}_{1j}\mathbf{u}_j||^2 + \tau^2\sum_{k=1}^{m_j}\mathbf{w}_{1jk}^\top\mathbf{w}_{1jk}v_{jk}^2}{2\sigma_e^2} \\
&+ \alpha_j\mathbf{u}_j^\top\widehat{\mathbf{S}}_j^{-2}\widehat{\boldsymbol{\gamma}}_j - \frac{1}{2}\alpha_j^2\mathbf{u}_j^\top\widehat{\mathbf{S}}_j^{-1}\widehat{\mathbf{R}}_j\widehat{\mathbf{S}}_j^{-1}\mathbf{u}_j - \frac{\alpha_j^2}{2}(\mathbf{v}_j\odot\mathbf{v}_j)^\top\mathrm{diag}(\widehat{\mathbf{S}}_j^{-1}\widehat{\mathbf{R}}_j\widehat{\mathbf{S}}_j^{-1}) \\
&- \frac{m_j}{2}\log(2\pi\sigma_{\gamma_j}^2) - \frac{||\mathbf{u}_j||^2 + ||\mathbf{v}_j||^2}{2\sigma_{\gamma_j}^2} + \frac{1}{2}\sum_{k=1}^{m_j}\log|2\pi e v_{jk}^2| + const.
\end{aligned}
\tag{S13}
$$

#### 2.3 The VB M-step

As we have obtained the analytical form of the ELBO, we can derive the updating formula of model parameters by setting the corresponding derivative equal to 0. The derivative with respective to $\sigma_e^2$, $\sigma_{\gamma_j}^2$ $\alpha_j$, $\boldsymbol{\beta}$ and $\tau$ are

$$
\begin{aligned}
\frac{\partial\mathcal{L}(q)}{\partial\sigma_e^2} &= -\frac{n_1}{2\sigma_e^2} + \frac{||\mathbf{y}_j^* - \tau\mathbf{W}_{1j}\mathbf{u}_j||^2 + \tau^2\sum_{k=1}^{m_j}\mathbf{w}_{1jk}^\top\mathbf{w}_{1jk}v_{jk}^2}{2\sigma_e^4} = 0, \\
\frac{\partial\mathcal{L}(q)}{\partial\sigma_{\gamma_j}^2} &= -\frac{m_j}{2\sigma_{\gamma_j}^2} + \frac{||\mathbf{u}_j||^2 + ||\mathbf{v}_j||^2}{2\sigma_{\gamma_j}^4} = 0, \\
\frac{\partial\mathcal{L}(q)}{\partial\alpha_j} &= \mathbf{u}_j^\top\widehat{\mathbf{S}}_j^{-2}\widehat{\boldsymbol{\gamma}}_j - \alpha_j\mathbf{u}_j^\top\widehat{\mathbf{S}}_j^{-1}\widehat{\mathbf{R}}_j\widehat{\mathbf{S}}_j^{-1}\mathbf{u}_j - \alpha_j(\mathbf{v}_j\odot\mathbf{v}_j)^\top\mathrm{diag}(\widehat{\mathbf{S}}_j^{-1}\widehat{\mathbf{R}}_j\widehat{\mathbf{S}}_j^{-1}) = 0, \\
\frac{\partial\mathcal{L}(q)}{\partial\boldsymbol{\beta}} &= \frac{-\mathbf{X}^\top(\mathbf{y}_j - \mathbf{X}\boldsymbol{\beta} - \tau\mathbf{W}_{1j}\mathbf{u}_j)}{\sigma_e^2} = 0, \\
\frac{\partial\mathcal{L}(q)}{\partial\tau} &= \frac{-(\mathbf{W}_{1j}\mathbf{u}_j)^\top(\mathbf{y}_j^* - \tau\mathbf{W}_{1j}\mathbf{u}_j) + \tau\sum_{k=1}^{m_j}\mathbf{w}_{1jk}^\top\mathbf{w}_{1jk}v_{jk}^2}{\sigma_e^2} = 0.
\end{aligned}
\tag{S14}
$$

With some algebraic work, the updating equations for all parameters have the following forms,

$$\begin{aligned}
\sigma_e^2 &= \frac{||\mathbf{y}_j^* - \tau \mathbf{W}_{1j}\mathbf{u}_j||^2 + \tau^2 \sum_{k=1}^{m_j} \mathbf{w}_{1jk}^\top \mathbf{w}_{1jk} v_{jk}^2}{n_1}, \\
\sigma_{\gamma_j}^2 &= \frac{||\mathbf{u}_j||^2 + ||\mathbf{v}_j||^2}{m_j}, \\
\alpha_j &= \frac{\mathbf{u}_j^\top \widehat{\mathbf{S}}_j^{-2} \widehat{\boldsymbol{\gamma}}_j}{\mathbf{u}_j^\top \widehat{\mathbf{S}}_j^{-1} \widehat{\mathbf{R}}_j \widehat{\mathbf{S}}_j^{-1} \mathbf{u}_j + (\mathbf{v}_j \odot \mathbf{v}_j)^\top \mathrm{diag}(\widehat{\mathbf{S}}_j^{-1} \widehat{\mathbf{R}}_j \widehat{\mathbf{S}}_j^{-1})}, \\
\boldsymbol{\beta} &= (\mathbf{X}^\top \mathbf{X})^{-1} \mathbf{X}^\top (\mathbf{y}_j - \tau \mathbf{W}_{1j}\mathbf{u}_j), \\
\tau &= \frac{\mathbf{u}_j^\top \mathbf{W}_{1j}^\top \mathbf{y}_j^*}{\mathbf{u}_j^\top \mathbf{W}_{1j}^\top \mathbf{W}_{1j} \mathbf{u}_j + \sum_{k=1}^{m_j} \mathbf{w}_{1jk}^\top \mathbf{w}_{1jk} v_{jk}^2}.
\end{aligned} \tag{S15}$$

The reduction steps are

$$\begin{aligned}
\alpha_j &= \frac{\alpha_j}{\tau}, \\
\sigma_{\gamma_j}^2 &= \tau^2 \sigma_{\gamma_j}^2.
\end{aligned} \tag{S16}$$

#### 2.4 PX-VBEM algorithm

Finally, we can summarize the overall algorithm as follows,

**Algorithm 1:** PX-VBEM for model CoMM-S$^2$MMs

1 *Initialization*: The parameters $(\sigma_{\gamma_j}^2, \sigma_e^2)$ are initialized using linear mixed model. Meanwhile, we set $\alpha_j = 0$, $\boldsymbol{\beta} = (\mathbf{X}^\top \mathbf{X})^{-1} \mathbf{X}^\top \mathbf{y}_j$ $\tau = 1$.
2 **repeat**
3 **E-step:** Given the $\tau = \tau^{(t)} = 1$ and other parameters updated in t-th iteration $\alpha_j = \alpha_j^{(t)}$, $\sigma_e^2 = (\sigma_e^{(t)})^2$, $\sigma_{\gamma_j}^2 = (\sigma_{\gamma_j}^{(t)})^2$, the posterior mean and variance of hidden variable $\boldsymbol{\gamma}_j$ is updated by the equation (S4)
4 **M-step:** The updating formula is derived by setting $\frac{\partial \mathcal{L}(q)}{\partial \boldsymbol{\theta}} = 0$ and referring to equation (10)
5 **Reduction-step:** Rescale the parameters $\alpha_j^{(t+1)} = \frac{\alpha_j^{(t+1)}}{\tau^{(t+1)}}$, $(\sigma_{\gamma_j}^{(t+1)})^2 = (\tau^{(t+1)})^2 (\sigma_{\gamma_j}^{(t+1)})^2$ and reset $\tau^{(t+1)} = 1$
6 **until** *Converge*;

### 3 Details for S-PrediXcan

Here we briefly introduce the procedures of implementing S-PrediXcan [2], which is a summary-based method to detect the association between genes and complex traits. In essence, S-PrediXcan is a Wald test implemented by summary-level GWAS dataset. In general, it consists of two stages. In the first stage, similar to PrediXcan, it requires to calculate the predicted gene expression using the transcriptome dataset. The difference between S-PrediXcan:Ridge and S-PrediXcan:Enet lies the model we trained. For S-PrediXcan:Ridge, we adopt ridge regression for prediction while we use elastic net regression for S-PrediXcan:Enet. In the second stage, it calculates the Z-score of the association between genes and traits according to the following formula.

$$Z_j \approx \sum_{k \in Model_j} \omega_{kj} \frac{\widehat{\sigma}_k}{\widehat{\sigma}_j} \frac{\widehat{\beta}_k}{se(\widehat{\beta}_k)} \tag{S17}$$

where $\omega_{kj}$ is the weight for $k$-th SNP in $j$-th gene estimated in the first stage. Additionally, $\widehat{\beta}_k$ and $se(\widehat{\beta}_k)$ corresponds to the coefficients and standard deviation of $k$-th SNP in $j$-th gene from GWAS summary data. The definition of $\widehat{\sigma}_k$ and $\widehat{\sigma}_j$ is as follows

$$\begin{aligned} \widehat{\sigma}_k &= \sqrt{\widehat{Var}(\mathbf{w}_{jk}^{\text{ref}})} \\ \widehat{\sigma}_j &= \sqrt{\widehat{Var}(\sum_{k \in \text{Model}_j} \omega_{kj} \mathbf{w}_{jk}^{\text{ref}})} \end{aligned} \tag{S18}$$

Obviously, $\widehat{\sigma}_k$ and $\widehat{\sigma}_j$ denotes the standard deviations of $k$-th SNP in $j$-th gene and the predicted gene expression of $j$-th gene respectively. Note that we use the approximation in (S17) instead of equality because we ignore the factor close to one in GWAS context.

In the simulation and real data analysis, we implemented S-PrediXcan in R for the easy implementation and fast computation. In the first stage, we trained the ridge regression model ($\alpha = 0$) or elastic net regression model ($\alpha = 0.5$) using the package glmnet from R to obtain the predicted gene expression and weights for each SNP. In the second stage, we calculate the Z-score of $j$-th gene to assess the signifcance of the association according to the expression (S17).

### 4 Simulations

#### 4.1 Comparisons of the calibrated ELBO with the marginal log-likelihood

In this simulation study, we compare the calibrated ELBO from the PX-VBEM algorithm for model (4) with the marginal log-likelihood from the EM algorithm. To make the comparison feasible, we used the genotype data $\mathbf{W}_2$ as reference panel, in other words, the genotype data from GWAS was served as reference data. Other settings are the same as the one described in Section 4. Specifically, we chose $n_1 = 400$, $n_2 = 5,000$, $m_j = 100$, $\rho = 0.2$, 0.5 or 0.8, $h_C^2 = 0.05$, $\pi = 0.2$ or 1, and $h_T^2 = 0$, 0.001, 0.002 or 0.003. The number of simulation replicate is 2,000. As shown in Figures S1 and S2, one can observe that the calibrated ELBO from PX-VBEM algorithm as described in Section 3.1 is essentially very close to that from EM algorithm.

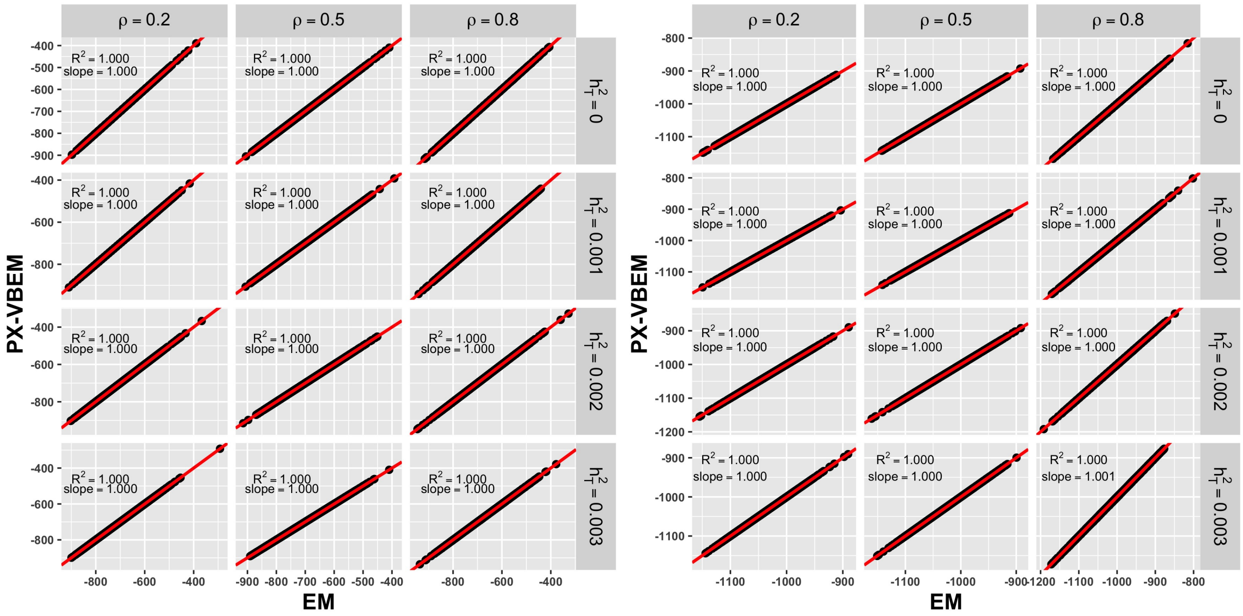

(a) $\pi = 0.2$ (b) $\pi = 1$

Figure S1: The scatter plot for the calibrated ELBO from PX-VBEM vs the marginal likelihood from EM with the setting $n_1 = 400$, $n_2 = 5{,}000$, $n_3 = 400$, $m_j = 100$ and $h_C^2 = 0.05$ for (a) $\pi = 0.2$ and (b) $\pi = 1$. The number of replication is 2000. The reference panel data is generated independently.

#### 4.2 Comparisons between CoMM-$S^2$ and CoMM

As the proposed approach, CoMM-$S^2$, is closely related to CoMM. The major difference is that CoMM-$S^2$ use GWAS summary statistics while CoMM use individual-level GWAS data. To better engage the performance of CoMM-$S^2$, we made side-side comparisons between CoMM-$S^2$ and CoMM using their corresponding test statistics. In this simulation study, we considered $n_1 = 400$, $n_2 = 5{,}000$, $n_3 = 400$, $m_j$=100, $\rho = 0.2$, 0.5 or 0.8, $\pi = 0.2$ or 1, and $h_T^2$= 0.001, 0.002 or 0.003. In addition, we consider two scenarios for reference panel, one from subsamples of GWAS data and the other from an independent data. For each combination of setting, we also provided the linear regression slope an $R^2$ of test statistics between CoMM-$S^2$ and CoMM. In all settings, the slope is close to 1 and $R^2$ is more than 97%. The plots for $\rho = 0.8$, $\pi = 1$ and subsamples in Figure S7 (b) can also be found in Figure 4 from the main text.

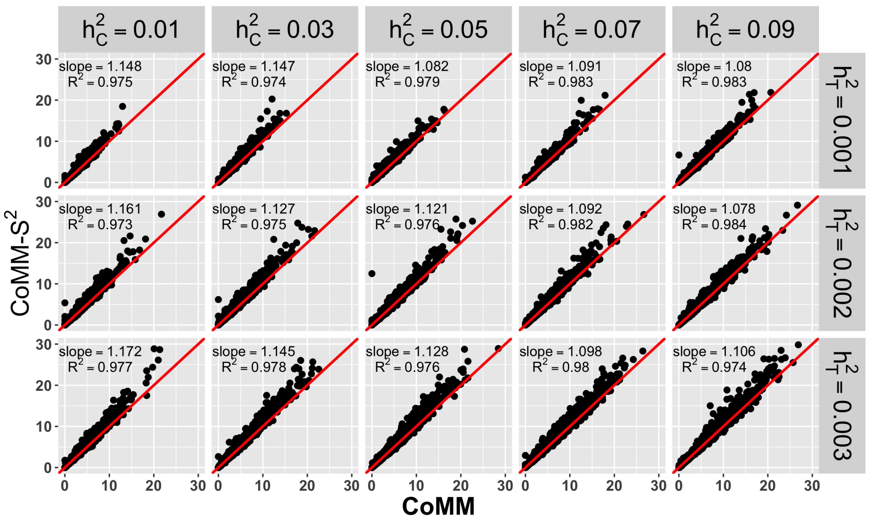

(a) Independent

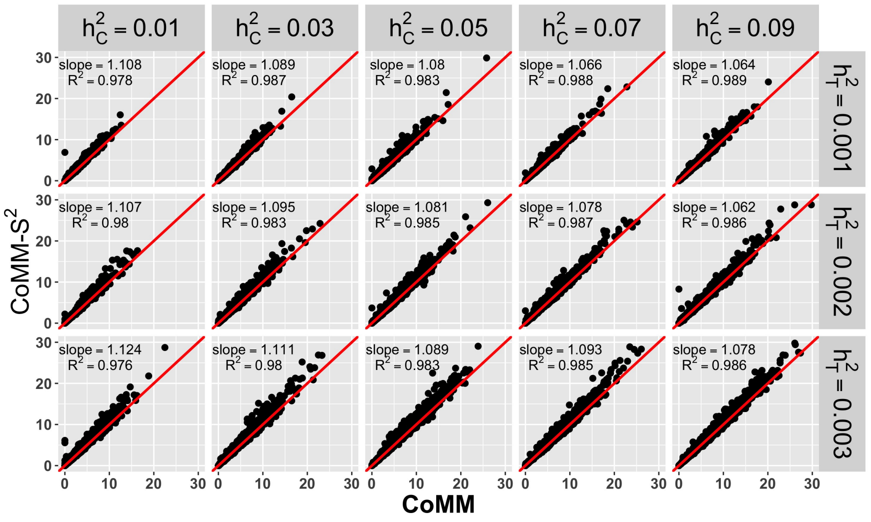

(b) Subsamples

Figure S2: The scatter plot of test statistics from LRT for CoMM-S$^2$ vs CoMM with the setting $n_1 = 400$, $n_2 = 5{,}000$, $n_3 = 400$, $m_j = 100$, $\rho = 0.2$, $\pi = 0.2$. The number of replication is 2000. The reference panel is generated from (a) an independent dataset and (b) subsamples from GWAS dataset.

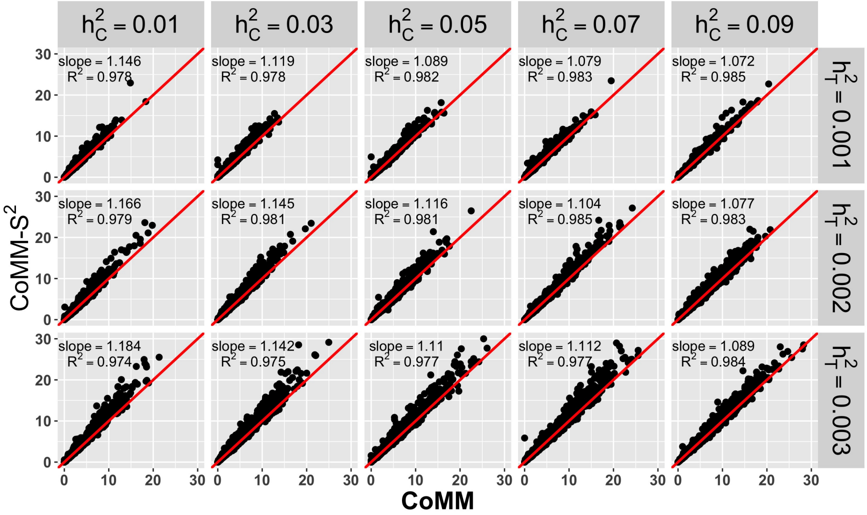

(a) Independent

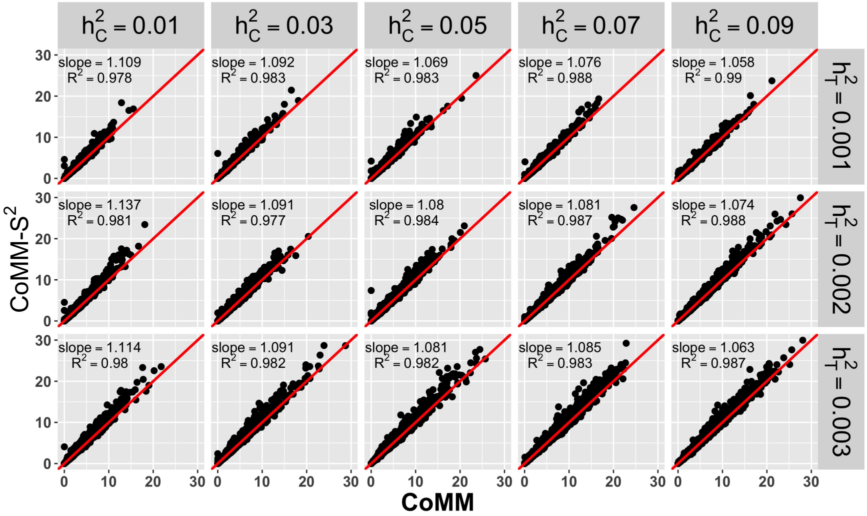

(b) Subsamples

Figure S3: The scatter plot of test statistics from LRT for CoMM-S$^2$ vs CoMM with the setting $n_1 = 400$, $n_2 = 5{,}000$, $n_3 = 400$, $m_j = 100$, $\rho = 0.2$, $\pi = 1$. The number of replication is 2000. The reference panel is generated from (a) an independent dataset and (b) subsamples from GWAS dataset.

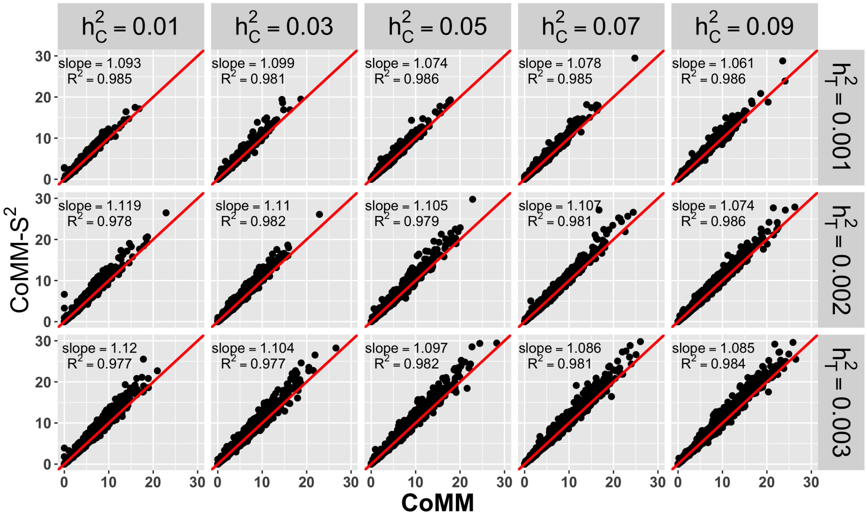

(a) Independent

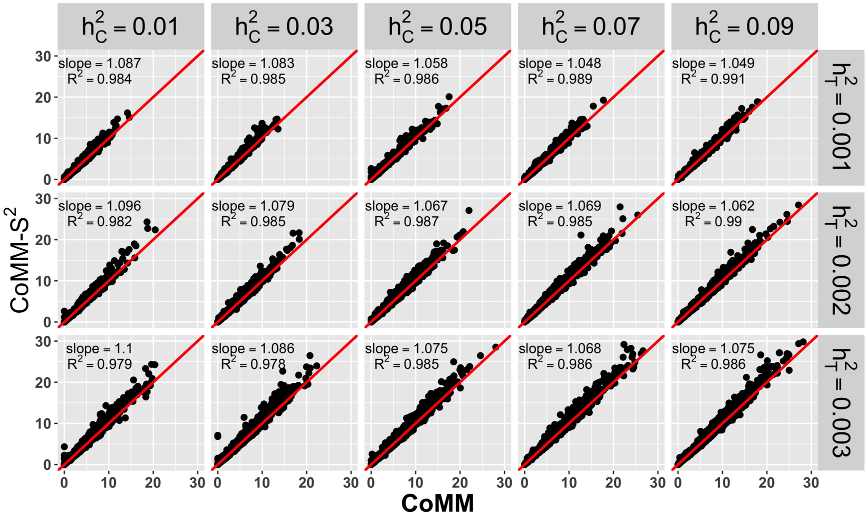

(b) Subsamples

Figure S4: The scatter plot of test statistics from LRT for CoMM-S$^2$ vs CoMM with the setting $n_1 = 400$, $n_2 = 5{,}000$, $n_3 = 400$, $m_j = 100$, $\rho = 0.5$, $\pi = 0.2$. The number of replication is 2000. The reference panel is generated from (a) an independent dataset and (b) subsamples from GWAS dataset.

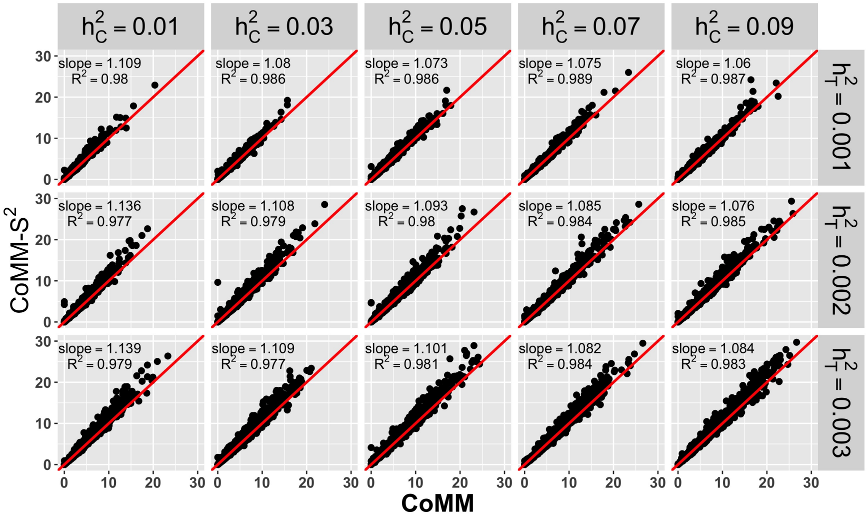

(a) Independent

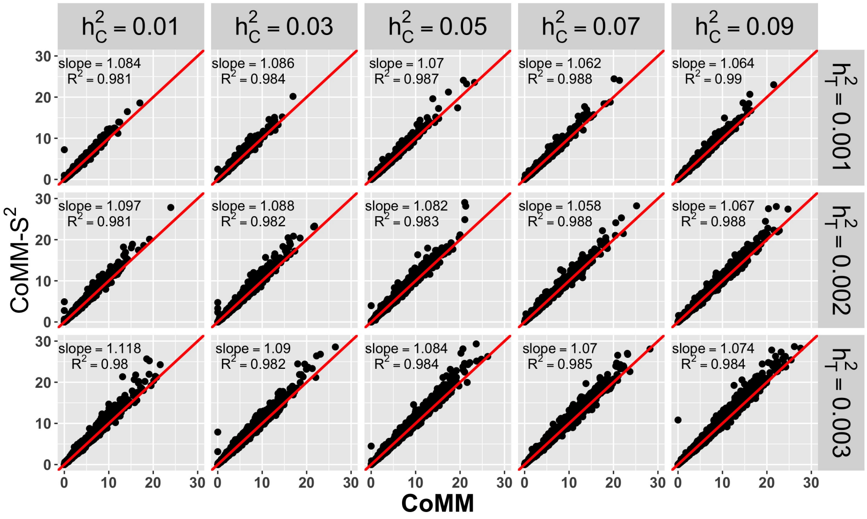

(b) Subsamples

Figure S5: The scatter plot of test statistics from LRT for CoMM-S$^2$ vs CoMM with the setting $n_1 = 400$, $n_2 = 5{,}000$, $n_3 = 400$, $m_j = 100$, $\rho = 0.5$, $\pi = 1$. The number of replication is 2000. The reference panel is generated from (a) an independent dataset and (b) subsamples from GWAS dataset.

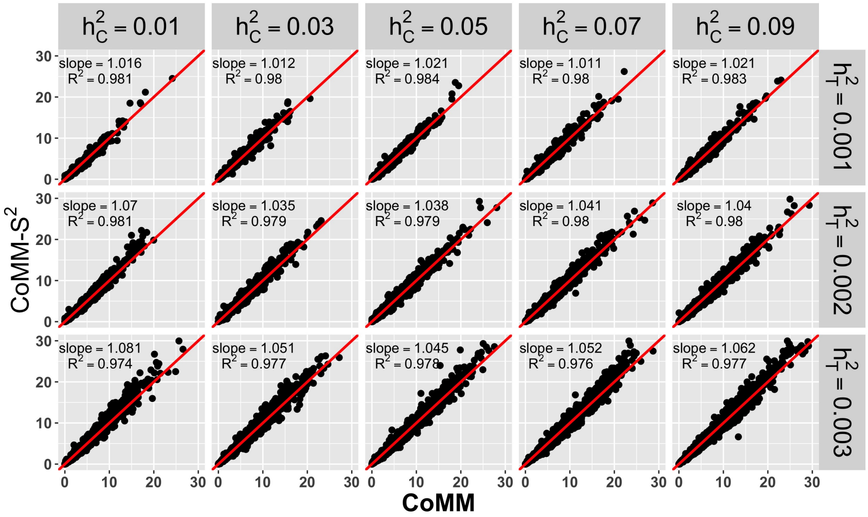

(a) Independent

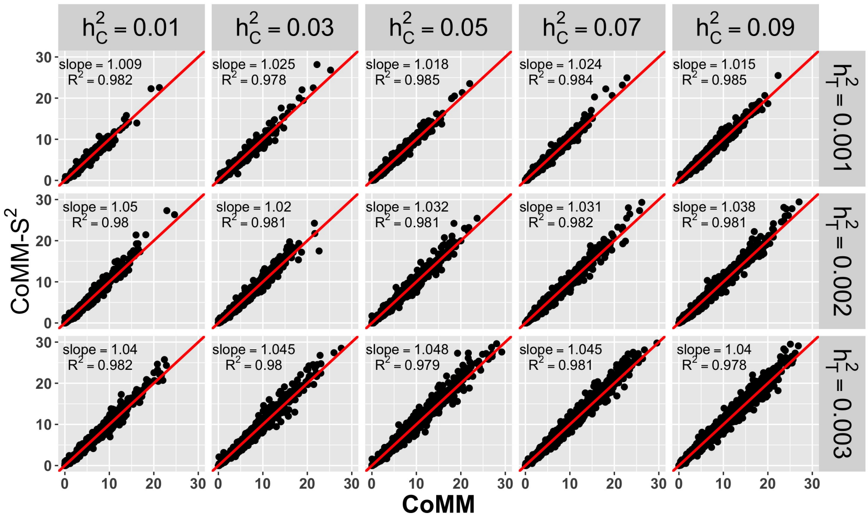

(b) Subsamples

Figure S6: The scatter plot of test statistics from LRT for CoMM-S$^2$ vs CoMM with the setting $n_1 = 400$, $n_2 = 5{,}000$, $n_3 = 400$, $m_j = 100$, $\rho = 0.8$, $\pi = 0.2$. The number of replication is 2000. The reference panel is generated from (a) an independent dataset and (b) subsamples from GWAS dataset.

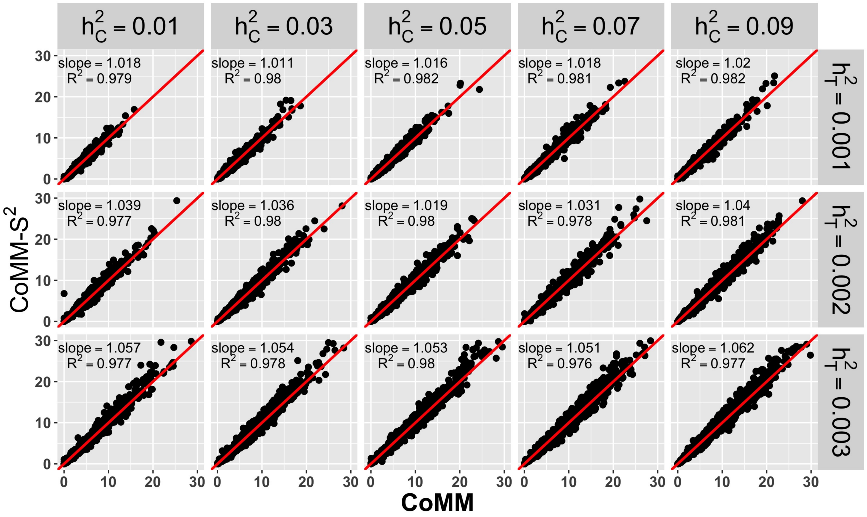

(a) Independent

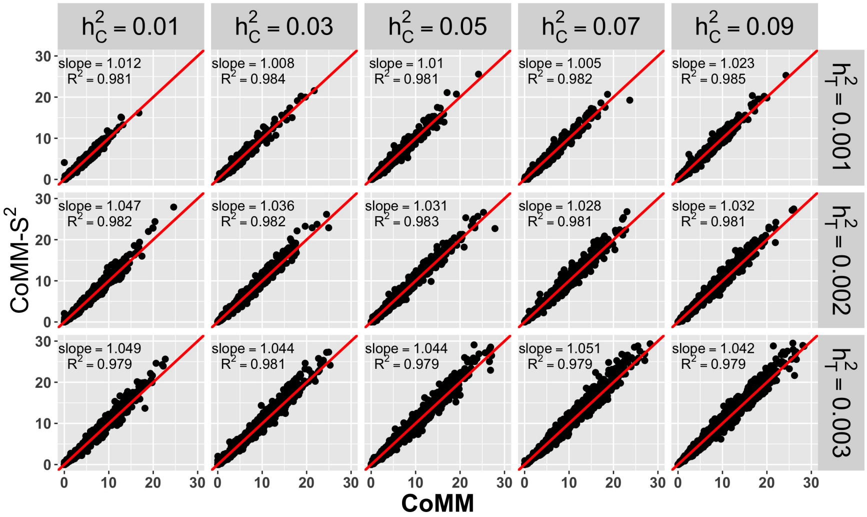

(b) Subsamples

Figure S7: The scatter plot of test statistics from LRT for CoMM-$S^2$ vs CoMM with the setting $n_1 = 400$, $n_2 = 5{,}000$, $n_3 = 400$, $m_j = 100$, $\rho = 0.8$, $\pi = 1$. The number of replication is 2000. The reference panel is generated from (a) an independent dataset and (b) subsamples from GWAS dataset.

##### 4.3 Comparisons of RL-SKAT with other methods

In the analysis of real data for NFBC1966 dataset and summary statistics for 15 traits, we observe that a tiny proportion of test statistics from CoMM degenerate to 0. It was demonstrated that under some regularity conditions, test statistics for LRT for the variance components follows a 50:50 mixture of $\chi^2_{\text{df}=0}$ and $\chi^2_{\text{df}=1}$ [18]. As shown in equation (14) of CoMM [19], CoMM tests for the null that $\alpha_j = 0$ which is equivalent to test both the mean and variance. We postulate that the degeneration of a small proportion of test statistics could be primarily due to the LRT involving the variance. To better understand this phenomenon, we conducted simulations to compare two-sample methods, *i.e.* CoMM-S$^2$, CoMM, S-Predixcan, and Predixcan, with RL-SKAT in the case that $h^2_C = 0$. Specifically, we fixed $n_1 = 400$, $n_2 = 5{,}000$, $n_3 = 400$, $m_j = 100$, $h^2_C = 0$, and $\pi$=0.2, and varied $\rho$= 0.2, 0.5 or 0.8, and $h^2_T = 0$ or 0.002. As test statistics from RL-SKAT follows a weighted mixture of chi-square distribution, we obtained its $p$-values and then transformed them to test statistics with one degree of freedom.

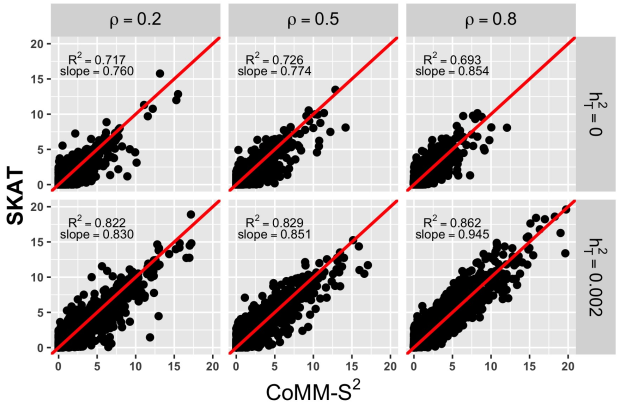

(a) RL-SKAT vs CoMM-S$^2$

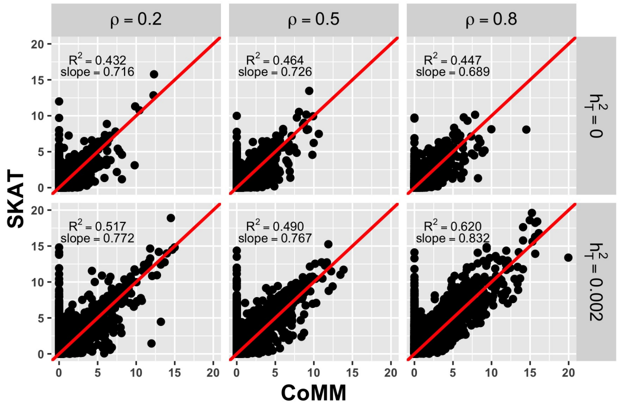

(b) RL-SKAT vs CoMM

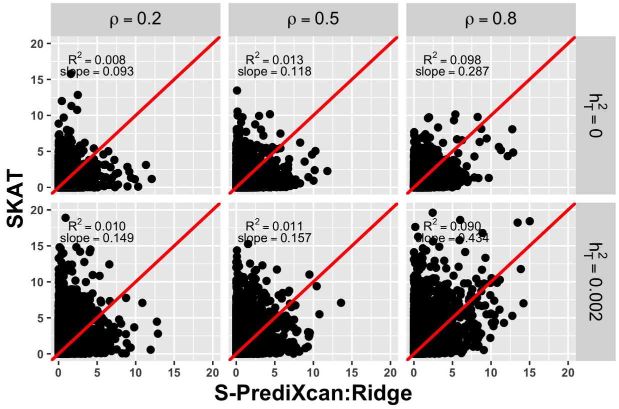

(c) RL-SKAT vs S-PrediXcan:Ridge

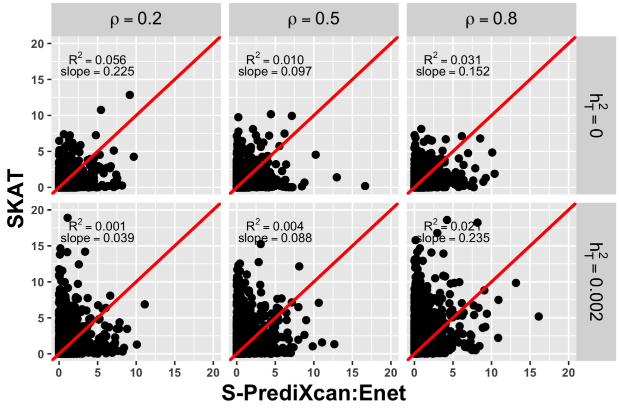

(d) RL-SKAT vs S-PrediXcan:Enet

Figure S8: The scatter plot of LRT statistics for SKAT vs other methods, (a) CoMM-S$^2$, (b) CoMM, (c) S-PrediXcan:Ridge and (d) S-PrediXcan:Enet, with the setting $n_1 = 400$, $n_2 =$ 5,000, $n_3 = 400$, $m_j = 100$, $h^2_C = 0$, $\pi = 0.2$. The number of replication is 2000.

#### 4.4 Comparisons for qq-plots under the null

In the null simulations, we examine the qq-plots for CoMM-S$^2$ together with other alternative methods (S-Predixcan:Ridge and S-Predixcan:Enet) under the null. In all settings, we chose $n_1 = 400$, $n_2 = 5{,}000$, and $n_3 = 400$ and varied $\rho \in \{0.2, 0.5, 0.8\}$, $h_T^2 \in \{0, 0.001, 0.002, 0.003\}$, and sparsity of eQTL as 0.1, 0.2 ,0.3, 0.4 and 0.5 or dense (sparsity = 1). The qq-plot for $\rho = 0.8$ with sparsity = 0.1 and 1 can be found in Figure 2 from the main text.

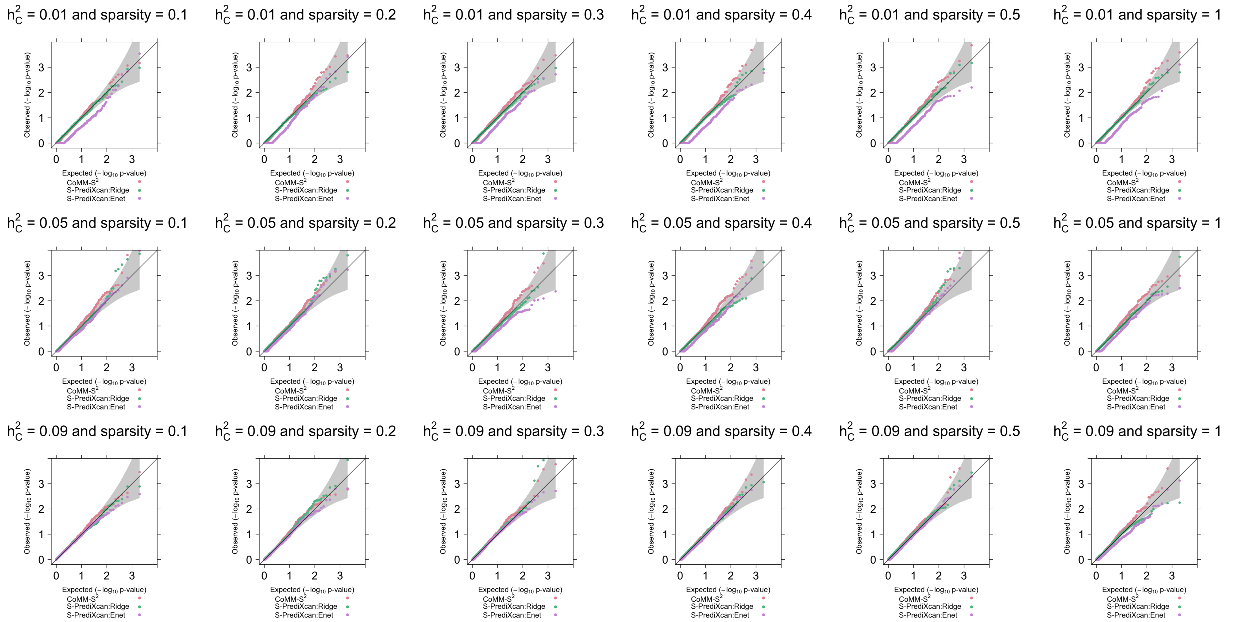

Figure S9: The qq-plot of $p$-values for each method (CoMM-S$^2$, S-Predixcan:Ridge and S-Predixcan:Enet) with the setting $n_1 = 400$, $n_2 = 5{,}000$, $n_3 = 400$, $\rho = 0.2$. The number of replication is 1000. From left to right, the sparsity varies $\pi \in \{0.1, 0.2, 0.3, 0.4, 0.5, 1\}$. From top to bottom, the cellular-level heritability varies from 0.01, 0.05 to 0.09.

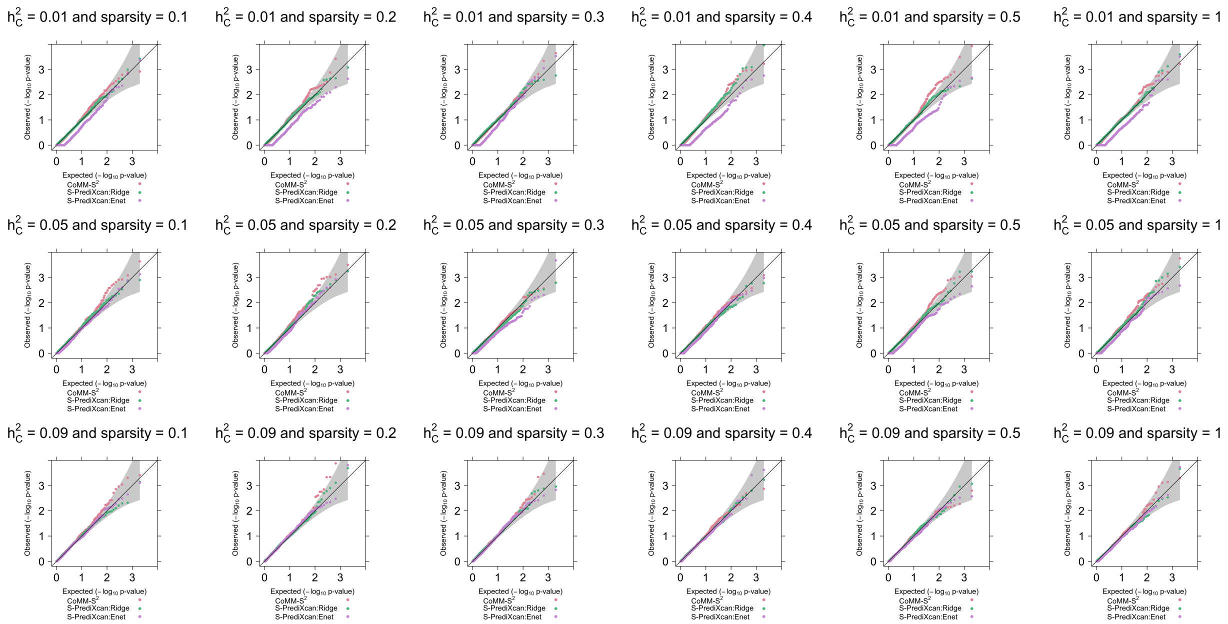

Figure S10: The qq-plot of $p$-values for each method (CoMM-S$^2$, S-Predixcan:Ridge and S-Predixcan:Enet) with the setting $n_1 = 400$, $n_2 = 5{,}000$, $n_3 = 400$, $\rho = 0.5$. The number of replication is 1000. From left to right, the sparsity varies $\pi \in \{0.1, 0.2, 0.3, 0.4, 0.5, 1\}$. From top to bottom, the cellular-level heritability varies from 0.01, 0.05 to 0.09.

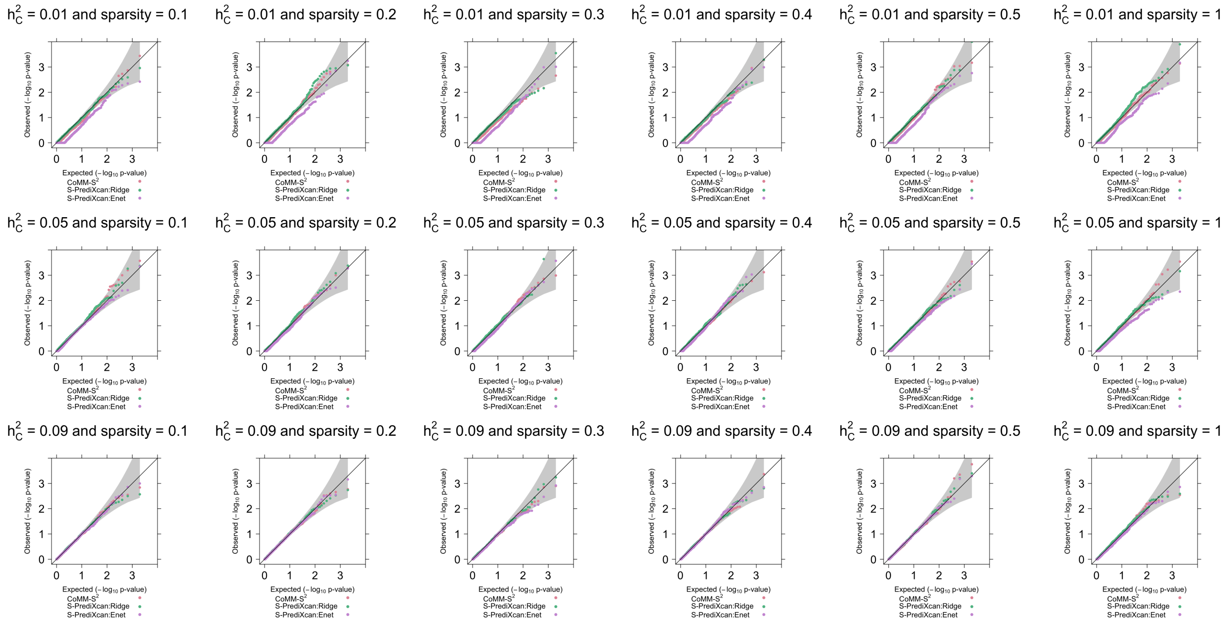

Figure S11: The qq-plot of $p$-values for each method (CoMM-S$^2$, S-Predixcan:Ridge and S-Predixcan:Enet) with the setting $n_1 = 400$, $n_2 = 5{,}000$, $n_3 = 400$, $\rho = 0.8$. The number of replication is 1000. From left to right, the sparsity varies $\pi \in \{0.1, 0.2, 0.3, 0.4, 0.5, 1\}$. From top to bottom, the cellular-level heritability varies from 0.01, 0.05 to 0.09.

#### 4.5 Comparison of power

In the power simulations, we examine the bar plots of power for CoMM-S$^2$ together with other alternative methods (S-Predixcan:Ridge and S-Predixcan:Enet) under the null. In all settings, we chose $n_1 = 400$, $n_2 = 5{,}000$, and $n_3 = 400$ and varied $\rho \in \{0.2, 0.5, 0.8\}$, $h_T^2 \in \{0, 0.001, 0.002, 0.003\}$, and sparsity of eQTL as 0.1, 0.2 ,0.3, 0.4 and 0.5 or dense (sparsity = 1). The plots for $\rho = 0.8$ can be found in Figure 3 from the main text.

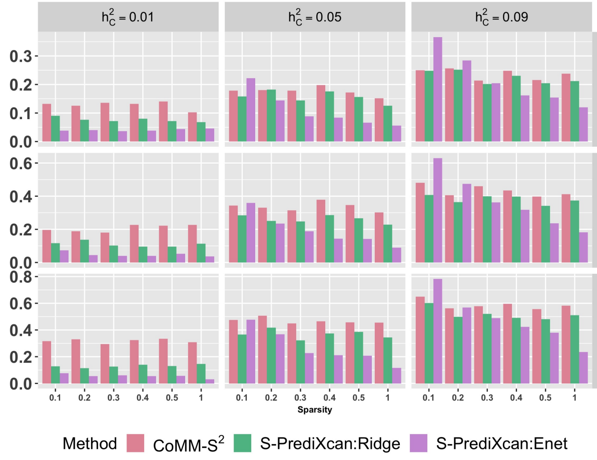

(a) $\rho = 0.2$

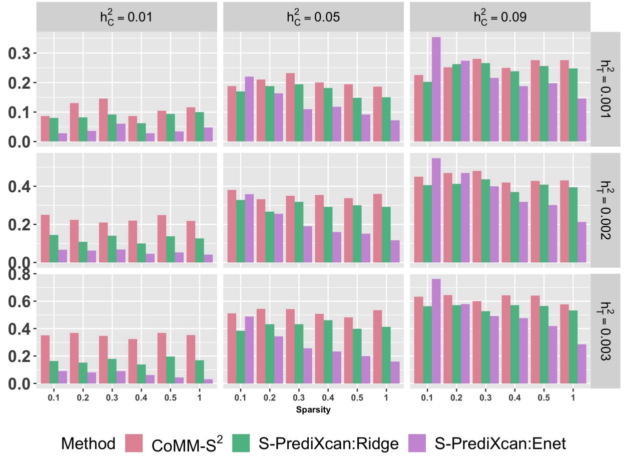

(b) $\rho = 0.5$

Figure S12: The comparison of power for CoMM-S$^2$, S-Predixcan:Ridge and S-Predixcan:Enet with the setting $n_1 = 400$, $n_2 = 5{,}000$, $n_3 = 400$, with (a) $\rho = 0.2$ and (b) $\rho = 0.5$. The number of replication is 500. For each subplot, the x-axis stands for the sparsity of SNP and the y-axis stands for the proportion of significant genes within 500 replications.

### 5 Real data analysis

#### 5.1 Comparisons between CoMM-S$^2$ and CoMM using NFBC1966 dataset

The scatter plot of test statistics of LRT for CoMM against CoMM-S$^2$ using 1KG samples as reference data. Note that we removed genes with cellular heritability less than 0.001 as the test statistics for tiny cellular heritability using CoMM is not reliable as shown in Figure S8 (b). One can observe that test statistics from CoMM-S$^2$ in the null region match these from CoMM well but inflated in the non-null region. As discussed in the main text, this is primarily due to use the 1KG as reference panel and Finnish samples from NFBC1966 dataset are distinct from European populations. For genes with cellular heritability less than 0.001, we evaluated the test statistics from RL-SKAT and made contrast for CoMM-S$^2$ or CoMM with RL-SKAT as shown in Figure S14. In comparison, we considered reference panel from both 400 subsamples and 1KG as shown in Figure S14 (a) and (b), respectively. One can observe the phenomenon that a certain proportion of test statistics from CoMM degenerates to zero while CoMM-S$^2$ does not. In addition, we notice that a few number of slopes in these scatter plots are less than 1, which is also consistent with simulation studies of Yang et al. [19] that RL-SKAT is more powerful in the region of small cellular heritability. Again, test statistics from RL-SKAT was calculated by transforming its $p$-values to chi-square distribution with one degree of freedom.

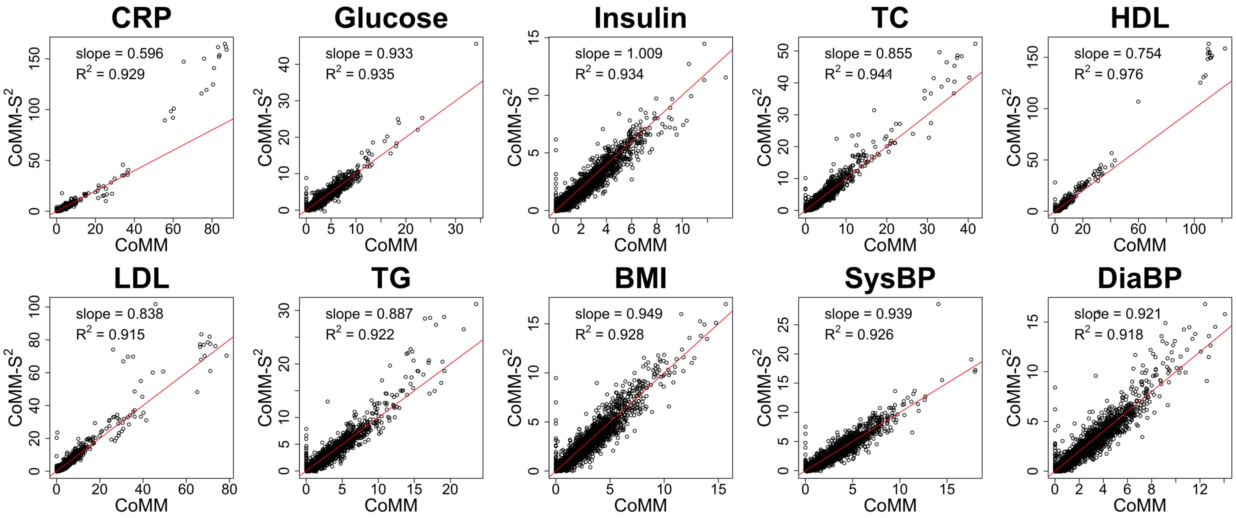

Figure S13: The scatter plot of test statistics from LRT for CoMM-S$^2$ vs CoMM, with transcriptiome data GTEx tissue Adipose Subcutaneous, GWAS data NFBC ,reference panel data 1KG. We remove the points with cellular heritability less than 0.01

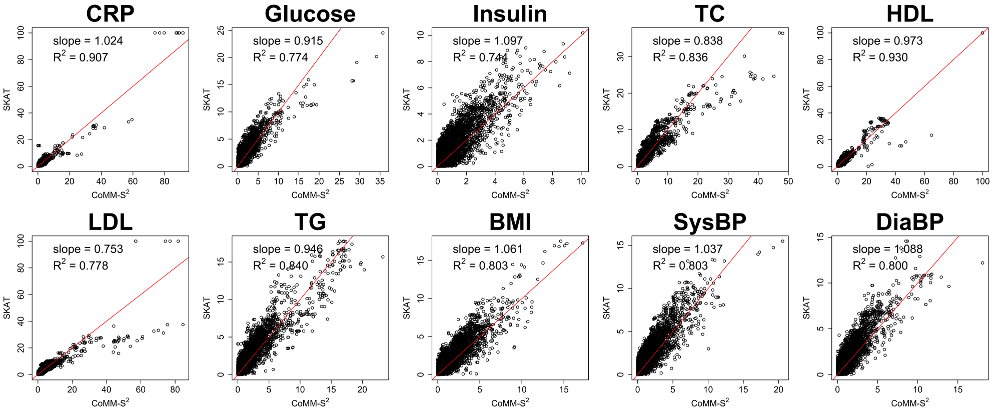

(a) CoMM-S$^2$ with 400 subsamples as reference panel.

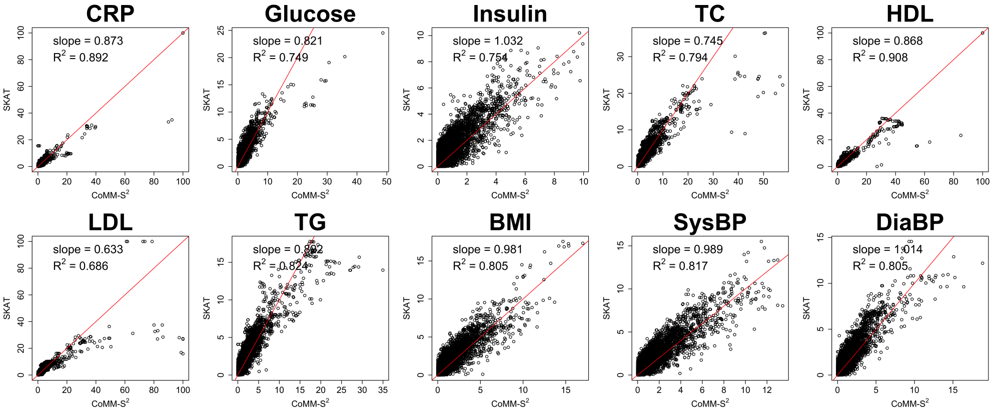

(b) CoMM-S$^2$ with 1KG as reference panel.

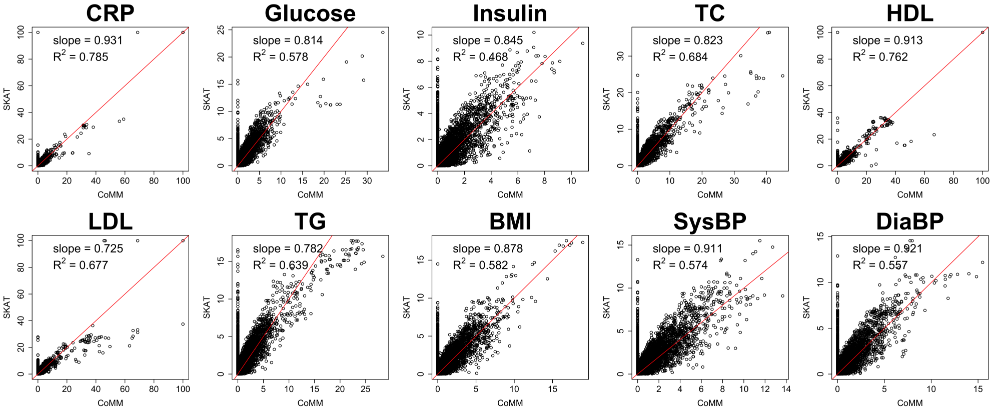

(c) CoMM using individual-level data.

Figure S14: The scatter plot of test statistics for CoMM-S$^2$ and CoMM vs RL-SKAT, respectively for genes having cellular heritability less than 0.01.

#### 5.2 The details for 14 traits

The details for 14 traits used to demonstrate the utility of CoMM-$S^2$ are give in Tables S1 and S2.

| | ID | Trait | Consortium |
|---|---|---|---|
| 1 | AD[13], | Alzheimer's disease | International Genomics of Alzheimer's Project(IGAP) |
| 2 | ASD | Autism spectrum disorder | Psychiatric Genomics Consortium (PGC) |
| 3 | CAD [14] | Coronary artery disease | Coronary ARtery DIsease Genome wide Replication and Meta-analysis (CARDIoGRAM) plus The Coronary Artery Disease (C4D) Genetics(CARDIoGRAMplusC4D) consortium |
| 4 | MI [4]. | myocardial infarction | UK Biobank cohort studies |
| 5 | SCZ | schizophrenia [10] | Psychiatric Genomics Consortium (PGC) |
| 6 | T2D [15] | Type 2 diabetes | DIAbetes Genetics Replication And Meta-analysis (DIAGRAM) consortium |
| 7 | BMIPA [6] | BMI for physically active individuals | The Genetic Investigation of ANthropometric Traits (GIANT) consortium |
| 8 | BMIPI [6] | BMI for physically inactive individuals | The Genetic Investigation of ANthropometric Traits (GIANT) consortium |
| 9 | BMIPAM [6] | BMI for physically active individuals in men | The Genetic Investigation of ANthropometric Traits (GIANT) consortium |
| 10 | BMIPAW [6] | BMI for physically active individuals in women | The Genetic Investigation of ANthropometric Traits (GIANT) consortium |
| 11 | BMIPIM [6] | BMI for physically inactive individuals in men | The Genetic Investigation of ANthropometric Traits (GIANT) consortium |
| 12 | BMIPIW [6] | BMI for physically inactive individuals in women | The Genetic Investigation of ANthropometric Traits (GIANT) consortium |
| 13 | BMIadjPAM [6] | BMI adjusted for physical activity for individuals in men | The Genetic Investigation of ANthropometric Traits (GIANT) consortium |
| 14 | BMIadjPAW [6] | BMI adjusted for physical activity for individuals in women | The Genetic Investigation of ANthropometric Traits (GIANT) consortium |

Table S1: The Consortia of fifteen traits

| | ID | YEAR | Website |
|---|---|---|---|
| 1 | AD | 2013 | http://web.pasteur-lille.fr/en/recherche/u744/igap/igap_download.php |
| 2 | ASD | 2015 | https://www.med.unc.edu/pgc/results-and-downloads |
| 3 | CAD | 2011 | http://www.cardiogramplusc4d.org |
| 4 | MI | 2012 | http://geneatlas.roslin.ed.ac.uk/downloads/?traits=313 |
| 5 | SCZ | 2013 | https://www.med.unc.edu/pgc/results-and-downloads |
| 6 | T2D | 2017 | http://diagram-consortium.org/downloads.html |
| 7 | BMIPA | 2017 | https://portals.broadinstitute.org/collaboration/giant/index.php/GIANT_consortium_data_files |
| 8 | BMIPI | 2017 | https://portals.broadinstitute.org/collaboration/giant/index.php/GIANT_consortium_data_files |
| 9 | BMIPAM | 2017 | https://portals.broadinstitute.org/collaboration/giant/index.php/GIANT_consortium_data_files |
| 10 | BMIPAW | 2017 | https://portals.broadinstitute.org/collaboration/giant/index.php/GIANT_consortium_data_files |
| 11 | BMIPIM | 2017 | https://portals.broadinstitute.org/collaboration/giant/index.php/GIANT_consortium_data_files |
| 12 | BMIPIW | 2017 | https://portals.broadinstitute.org/collaboration/giant/index.php/GIANT_consortium_data_files |
| 13 | BMIadjPAM | 2017 | https://portals.broadinstitute.org/collaboration/giant/index.php/GIANT_consortium_data_files |
| 14 | BMIadjPAW | 2017 | https://portals.broadinstitute.org/collaboration/giant/index.php/GIANT_consortium_data_files |

Table S2: The website of fifteen traits

#### 5.3 Analysis results for 14 traits

For the analysis of 14 traits using GTEx transcriptome data, we additionally report the total number of genes identified across all tissues for three methods, *i.e.* , CoMM-$S^2$, S-PrediXcan:Enet and S-PrediXcan:Ridge, in Table S3, and the corresponding qq-plot for these traits across all tissues is shown in Figure S15.

| | CoMM-$S^2$ | S-PrediXcan:Enet | S-PrediXcan:Ridge |
|---|---|---|---|
| AD | 4614(1103) | 398(165) | 664(284) |
| ASD | 2(0) | 1(0) | 3(0) |
| CAD | 1811(358) | 94(23) | 158(28) |
| MI | 2503(144) | 188(32) | 237(38) |
| SCZ | 214(47) | 69(19) | 79(12) |
| T2D | 4939(956) | 216(87) | 319(93) |
| BMIPA | 5405(1428) | 315(100) | 605(189) |
| BMIPI | 1107(337) | 27(10) | 64(18) |
| BMIPAM | 460(107) | 10(2) | 32(3) |
| BMIPAW | 1640(503) | 126(35) | 159(51) |
| BMIPIM | 287(96) | 6(2) | 4(2) |
| BMIPIW | 351(102) | 17(4) | 26(6) |
| BMIadjPAM | 2423(627) | 123(39) | 282(91) |
| BMIadjPAW | 4638(1226) | 210(80) | 343(114) |

Table S3: The number of significant genes identified across the tissues at the significant level ($5 \times 10^{-6}$) for each method. In this table, we simply aggregate all the identified genes across the tissues. The number in the parenthesis denoted the genes reported on the GWAS catalog.

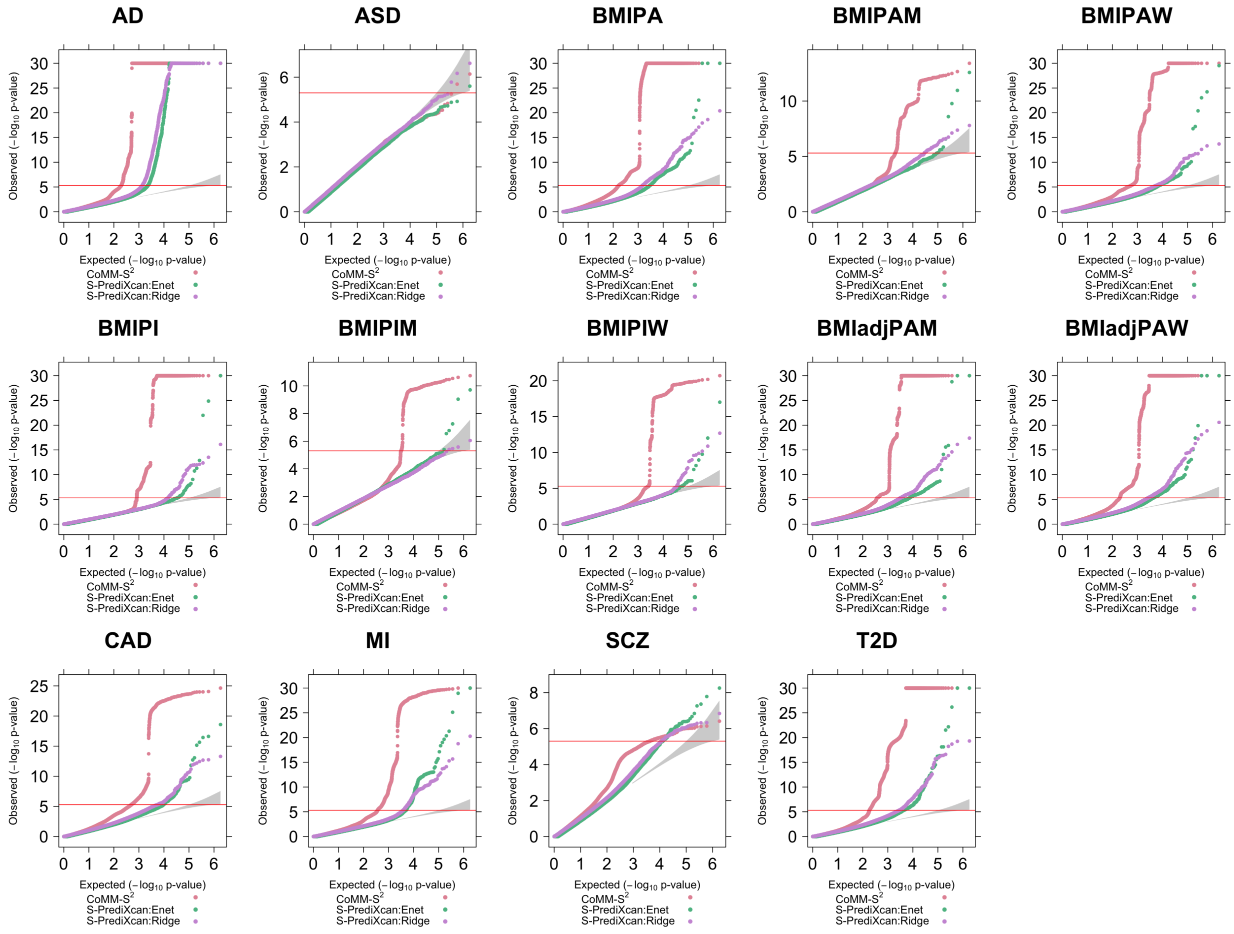

Figure S15: The qq-plot of $p$-values for 14 traits. 1KG is used as reference panel and GTEx data across 48 tissues are used as transcriptome data. For the qq-plot, we conducted genomic control for all three methods.

[14] Heribert Schunkert, Inke R König, Sekar Kathiresan, Muredach P Reilly, Themistocles L Assimes, Hilma Holm, Michael Preuss, Alexandre FR Stewart, Maja Barbalic, Christian

Gieger, et al. Large-scale association analysis identifies 13 new susceptibility loci for coronary artery disease. *Nature genetics*, 43(4):333, 2011.

[15] Robert A Scott, Laura J Scott, Reedik Mägi, Letizia Marullo, Kyle J Gaulton, Marika Kaakinen, Natalia Pervjakova, Tune H Pers, Andrew D Johnson, John D Eicher, et al. An expanded genome-wide association study of type 2 diabetes in europeans. *Diabetes*, 66(11):2888–2902, 2017.

[16] Oliver Stegle, Leopold Parts, Richard Durbin, and John Winn. A bayesian framework to account for complex non-genetic factors in gene expression levels greatly increases power in eqtl studies. *PLoS computational biology*, 6(5):e1000770, 2010.

[17] Oliver Stegle, Leopold Parts, Matias Piipari, John Winn, and Richard Durbin. Using probabilistic estimation of expression residuals (peer) to obtain increased power and interpretability of gene expression analyses. *Nature protocols*, 7(3):500, 2012.

[18] Daniel O Stram and Jae Won Lee. Variance components testing in the longitudinal mixed effects model. *Biometrics*, pages 1171–1177, 1994.

[19] Can Yang, Xiang Wan, Xinyi Lin, Mengjie Chen, Xiang Zhou, and Jin Liu. Comm: a collaborative mixed model to dissecting genetic contributions to complex traits by leveraging regulatory information. *Bioinformatics*, 2018.
